# PEARL-catalyzed peptide bond formation after chain reversal during the biosynthesis of non-ribosomal peptides

**DOI:** 10.1101/2023.12.23.573212

**Authors:** Yue Yu, Wilfred A. van der Donk

**Affiliations:** Department of Chemistry and Howard Hughes Medical Institute, University of Illinois at Urbana-Champaign, Urbana, IL 61801

## Abstract

A subset of nonribosomal peptide synthetases (NRPSs) and polyketide synthases (PKSs) are encoded in their biosynthetic gene clusters (BGCs) with enzymes annotated as lantibiotic dehydratases. The functions of these putative lantibiotic dehydratases remain unknown. Here, we characterize an NRPS-PKS BGC with a putative lantibiotic dehydratase from the bacterium *Stackebrandtia nassauensis* (*sna*). Heterologous expression revealed several metabolites produced by the BGC, and the omission of selected biosynthetic enzymes revealed the biosynthetic sequence towards these compounds. The putative lantibiotic dehydratase catalyzes peptide bond formation that extends the peptide scaffold opposite to the NRPS and PKS biosynthetic direction. The condensation domain of the NRPS catalyzes the formation of a ureido group, and bioinformatics analysis revealed distinct active site residues of ureido-generating condensation (UreaC) domains. This work demonstrates that the annotated lantibiotic dehydratase serves as a separate amide bond-forming machinery in addition to the NRPS, and that the lantibiotic dehydratase enzyme family possesses diverse catalytic activities in the biosynthesis of both ribosomal and non-ribosomal natural products.

## Introduction

Nonribosomal peptides (NRPs) are natural products that possess a range of biological activities, such as antibiotic,^1^ anticancer,^2^ biosurfactant,^3^ and immunosuppressant.^4^ Their peptide scaffold is biosynthesized by nonribosomal peptide synthetases (NRPSs), multimodular enzymes that work like assembly lines.^5-7^ A typical peptide elongation module consists of three domains: condensation (C), adenylation (A), and thiolation (T). A conserved serine of the T domain is posttranslationally modified by the addition of phosphopantetheine.^8,9^ The A domain activates an amino acid by adenylation and loads it onto the phosphopantetheine arm of the T domain as an acyl-thioester intermediate. The C domain then catalyzes the formation of peptide bonds between intermediates bound to the T domain to extend the peptide chain (Figure 1*A*).

**Figure 1.**
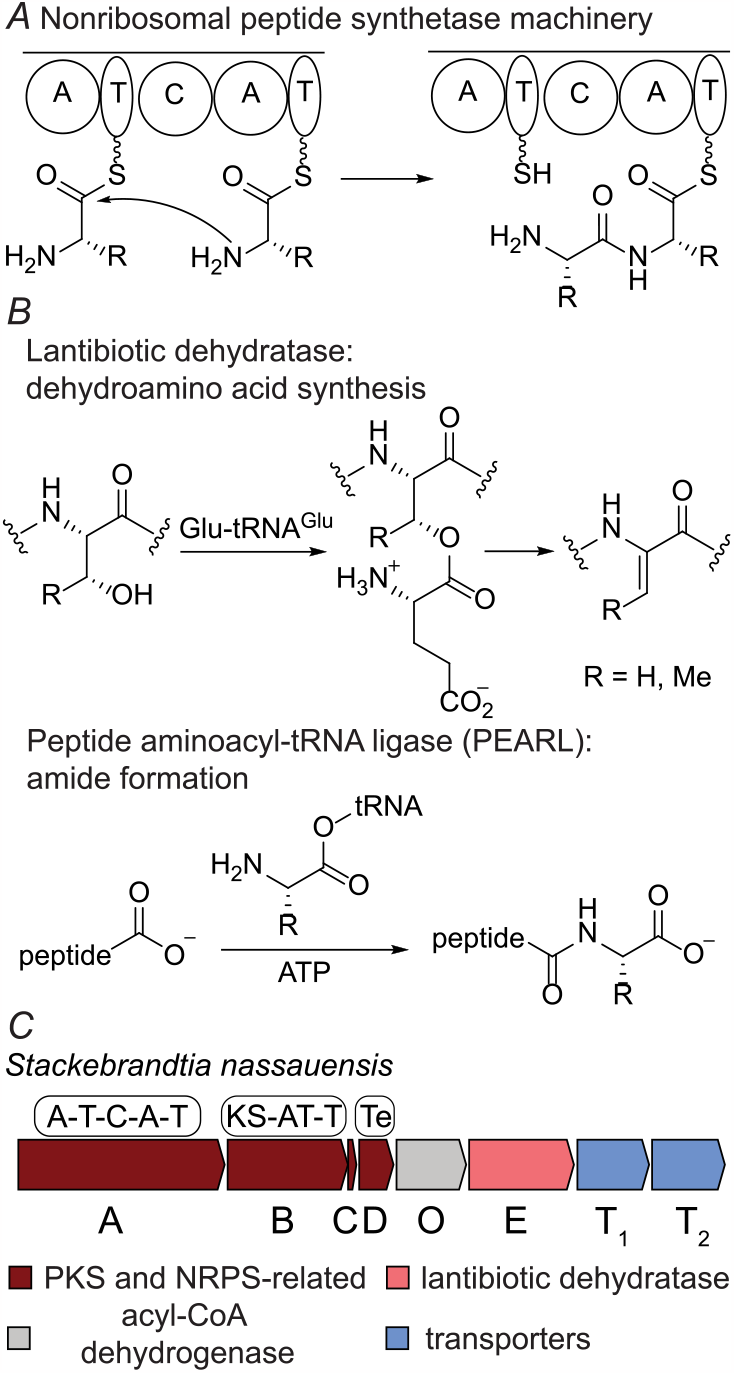
(*A*) The peptide bond formation chemistry of NRPSs. (*B*) Known enzymatic activities of the lantibiotic dehydratase enzyme family. (*C*) Schematic diagram of the *sna* BGC from *Stackebrandtia nassauensis*. KS: ketosynthase. Te: thioesterase. For the accession IDs for all proteins in the *sna* BGC, see the *Supporting Information*.

A subset of NRPSs are encoded in biosynthetic gene clusters (BGCs) with enzymes annotated as lantibiotic dehydratases ^10,11^ Lantibiotic dehydratases (protein family PF04738) generate the dehydroamino acids^12^ of ribosomally synthesized and post-translationally modified peptides (RiPPs),^13^ including lanthipeptides^11^ (called lantibiotics if they display antibiotic activity) and thiopeptides.^14^ The dehydration reaction involves the glutamylation of serine and threonine hydroxyl groups using glutamyl-tRNA^Glu^ and subsequent elimination of glutamate to generate peptidyl dehydroamino acids^10,15^ (Figure 1*B*). Other enzymes frequently mis-annotated as lantibiotic dehydratase are peptide aminoacyl-tRNA ligases (PEARLs).^16-18^ PEARLs catalyze peptide bond formation at the C-terminus of a carrier peptide using adenosine-5’-triphosphate (ATP) and aminoacyl-tRNA^17^ (Figure 1*B*). The amino acid added by the PEARL will undergo enzymatic modifications and proteolysis to yield amino acid-derived natural products.^19-21^ However, neither a RiPP precursor peptide nor a cognate PEARL carrier peptide can be identified in the NRPS BGCs, indicating the putative lantibiotic dehydratases serve a different function in the biosynthesis of NRPs.

In this study, we investigated a hybrid NRPS-PKS BGC^22-24^ from *Stackebrandtia nassauensis* that contains a putative lantibiotic dehydratase (Fig. 1*C*). Heterologous expression, comparative metabolomics, and structural elucidation revealed a series of novel metabolites. The biosynthetic sequence was revealed by omitting select biosynthetic enzymes during heterologous expression. The NRPS SnaA links two arginine amine groups through a ureido group, leaving an inert carboxylate at the initiation position that cannot be further extended by the NRPS machinery. The putative lantibiotic dehydratase SnaE catalyzes peptide bond formation at this unactivated carboxylate of the terminal ureido group, achieving chain extension in the opposite direction to NRPS-PKS biosynthesis. The results show that the annotated lantibiotic dehydratases that colocalize with NRPS/PKSs likely biosynthesize amide bonds that are not amenable to thioester assembly line biochemistry.

Ureido group formation is one of the many versatile reactions during NRP biosynthesis.^25,26^ In vitro studies of SylC in syringolin biosynthesis suggest that the ureido moiety likely originates from bicarbonate.^27^ This unusual head-to-head condensation reaction between two amino acids led us to hypothesize that the condensation domain of the NRPS SnaA is specialized for ureido group formation. Analysis of the condensation domain sequences associated with ureido-containing NRPs predicts that the active site signature of ureido-generating condensation (UreaC) domain is EHHXXHDG (X represents any amino acid) compared to the canonical XHHXXXDG motif for peptide bond formation.^9,28^ Condensation domains that do not generate ureido groups in the Minimum Information about a Biosynthetic Gene cluster (MiBiG) database^29^ never have the EHHXXHDG motif, suggesting the extra conservation of glutamate and histidine residues in the active site of C domains marks the signature of ureido group formation.

## Results

### Products produced by the *sna* BGC

Around two thousand NRPS/PKS BGCs contain enzymes annotated as lantibiotic dehydratase (NCBI, June 2023). Most of the annotated lantibiotic dehydratases are stand-alone enzymes, but some of them are fused to thioesterase or condensation domains. No lanthipeptide precursor peptides can be bioinformatically identified in these BGCs. This observation suggests that these putative lantibiotic dehydratases are involved in NRP or polyketide (PK) biosynthesis rather than lanthipeptide biosynthesis.

The BGCs in question are mostly from Actinobacteria. We chose one representative candidate gene cluster from *S. nassauensis* (Figure 1*C*) for heterologous expression in *Streptomyces albus* J1074.^*30*^ Expression of a construct containing *snaABCDOET*_*1*_*T*_*2*_ under control of the SP44 constitutive promoter^31^ produced several new metabolites detected by liquid chromatography-mass spectrometry (LC-MS). Expression in two liters of media, purification, and characterization by nuclear magnetic resonance spectroscopy revealed the structures of three major metabolites (compounds A, B, and C, Figure 2).

**Figure 2.**
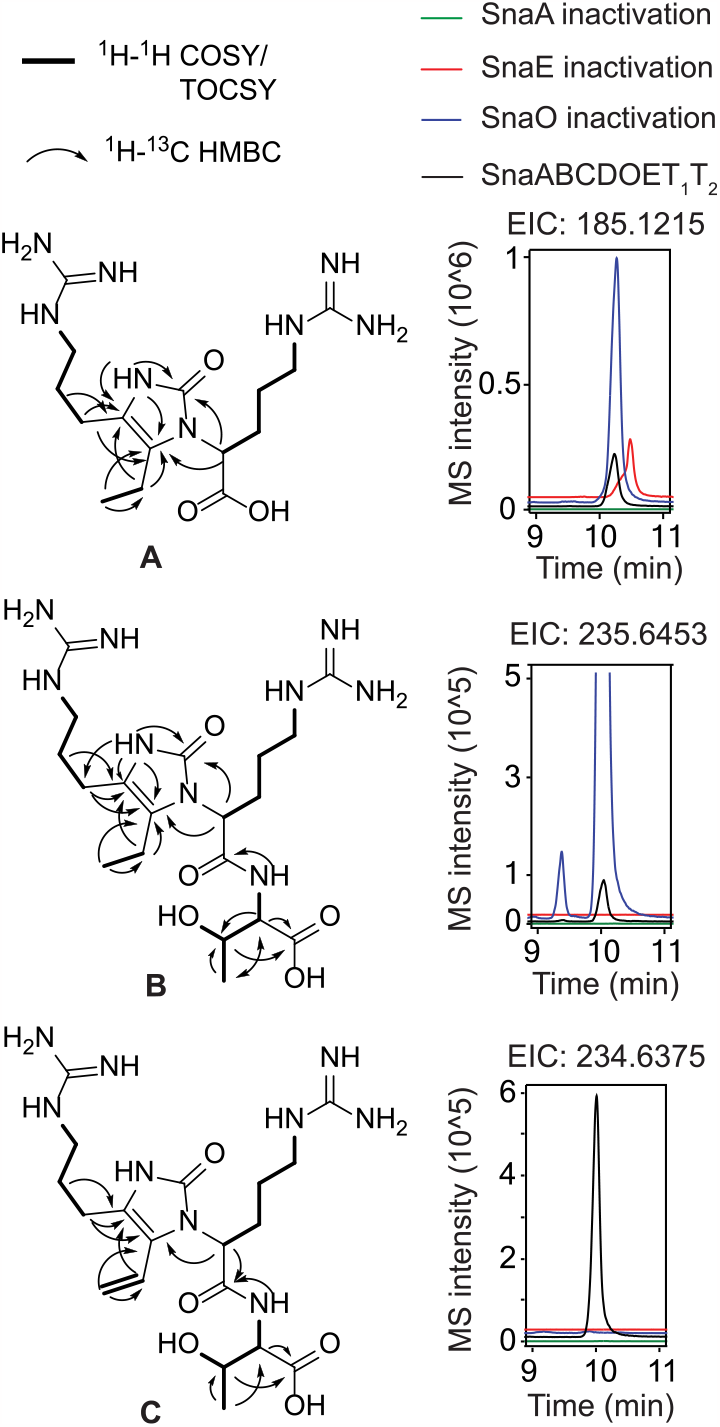
Structures of metabolites produced from the *sna* BGC using different heterologous expression constructs. EIC: extracted ion chromatogram. Only key NMR connectivities used to solve the structures are shown. For complete NMR data, see the Supporting Information.

To investigate the enzymes required to produce these compounds, *snaA* (NRPS), *snaO* (dehydrogenase), and *snaE* (putative lantibiotic dehydratase) were individually inactivated during heterologous expression (Figure 2). For *snaA*, both serine codons of the T domains were mutated to alanine to yield an inactive mutant that cannot be converted to the holo form. In-frame deletions were used to inactivate *snaO* and *snaE* (*Supporting Information*). Production of compound A with two arginines required SnaA but not SnaO or SnaE (Figure 2). Compound B contains one more Thr than compound A, and its biosynthesis required SnaA and SnaE but not SnaO. Compound C has an additional alkene group compared to compound B, and required SnaA, SnaE, and SnaO for biosynthesis. These results suggest that compound A is an early-stage biosynthetic intermediate produced by the NRPS and PKS (vide infra), and compound C is likely a later intermediate or the final product. Structural comparison between A and B strongly suggests that the putative lantibiotic dehydratase SnaE catalyzes the formation of a peptide bond between a threonine donor and a motif made by the NRPS/PKS. Therefore, SnaE is a peptide bond-forming enzyme rather than a dehydratase.

In addition to compounds A-C, we observed three other products, compounds D-F. Compounds A and D, B and E, and C and F are always produced together, respectively (Figure 2, 3*A*). High-resolution mass spectrometry suggests the difference in molecular formulae of each pair is H_2_O. A spontaneous intramolecular dehydrative cyclization between the ureido NH and the ketone explains the formation of compounds A, B, and C from D, E, and F, respectively. Similar reactions of guanidino nitrogens spontaneously cyclizing onto an arginine ethyl ketone have been observed in the study of saxitoxin biosynthesis.^32-34^ Compounds D-F eluded spectroscopic characterization because of their high cyclization reactivity during purification efforts, but high-resolution MS/MS spectra (Figure S1-3) as well as observed non-enzymatic conversion of D to A, E to B, and F to C during purification strongly support the structural assignment.

**Figure 3.**
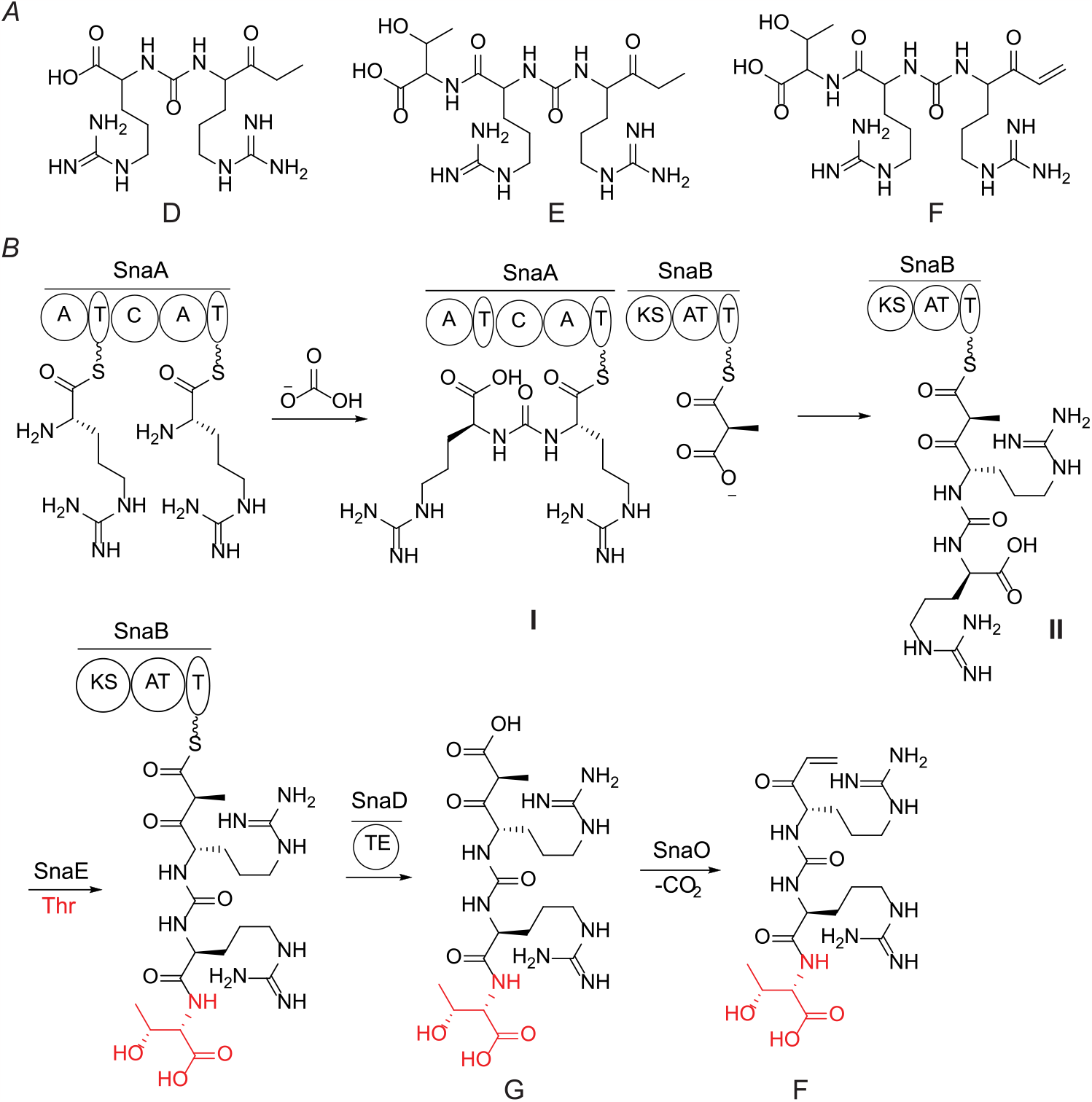
(*A*) Proposed structures of compounds D, E, and F. Stereochemistry could not be determined because of the high cyclization reactivity of these compounds. (*B*) The proposed biosynthetic sequence to generate compound F (threopeptin).

### Proposed biosynthetic pathway

Knowing the required enzymes for the biosynthesis of each metabolite, we propose the following biosynthetic sequence (Figure 3*B*). The two adenylation domains of SnaA (NCBI ADD43706.1) both activate and load arginine onto the peptidyl carrier protein (PCP) as activated thioesters. The condensation domain is unusual from bioinformatic analysis (vide infra) and catalyzes the condensation between two amine groups of arginine to form a ureido group that is likely derived from bicarbonate (HCO_3_^-^).^27^ The PKS SnaB (NCBI ADD43707.1) incorporates a propionate extension unit into the growing chain, as shown by isotope enrichment upon feeding 2-^13^C sodium propionate to the heterologous expression system (Figure S4). The SnaB-bound intermediate may be hydrolyzed by the thioesterase SnaD (NCBI ADD43709.1) to form compound D which upon cyclization gives compound A (Figure S5A). Threonine addition by SnaE (NCBI ADD43711.1) can occur on PCP/acyl carrier protein (ACP)-bound intermediates **I** or **II** (Figure 3*B*) or on the free molecules D or H (Figure S5). Based on precedence with the dehydrogenase EpnF,^35,36^ compound F could be produced from compound G by SnaO (NCBI ADD43710.1) via a decarboxylation-dehydrogenation reaction sequence, but conversion of compound E to F by SnaO cannot be ruled out. As outlined in the Discussion section, we consider compound F the final product of the pathway and term this compound threopeptin, whereas the formation of compounds A-E are proposed to be off-pathway via non-enzymatic cyclization and/or premature thioesterase activity (Figure S5).

### Biochemical and bioinformatic studies on ureido group formation

Ureido group formation is one of the many versatile reactions catalyzed by C domains during NRP biosynthesis.^25,26^ Based on current understanding, we could not have predicted that the *sna* BGC would produce a ureido structure. Therefore, we bioinformatically investigated whether the C domains associated with known ureido-containing natural products (UreaC domains) have a distinct active site amino acid signature. We compiled the UreaC domains in the MiBiG database based on collinearity to product structures (e.g. anabaenopeptins,^37^ bulbfieramide,^38^ chitinimide,^39^ and pseudovibriomide^40^) as well as examples with in vitro confirmation of the ureido formation enzyme activity (e.g. syringolin A,^27^ pacidamycin,^41^ antipain,^42^ and muraymycin^42^). Multiple sequence alignment showed that UreaC domains have a conserved EHHXXHDG active site (Figure 4*A*) compared to the canonical XHHXXXDG active site of the amide bond-forming C domains.^9,28^ Examining all C domain sequences in MiBiG showed that none of the C domains with other functions have an EHHXXHDG active site. Therefore, based on current examples, the EHHXXHDG signature of the C domain active site appears to be sufficient and necessary to indicate the ureido formation activity.

**Figure 4.**
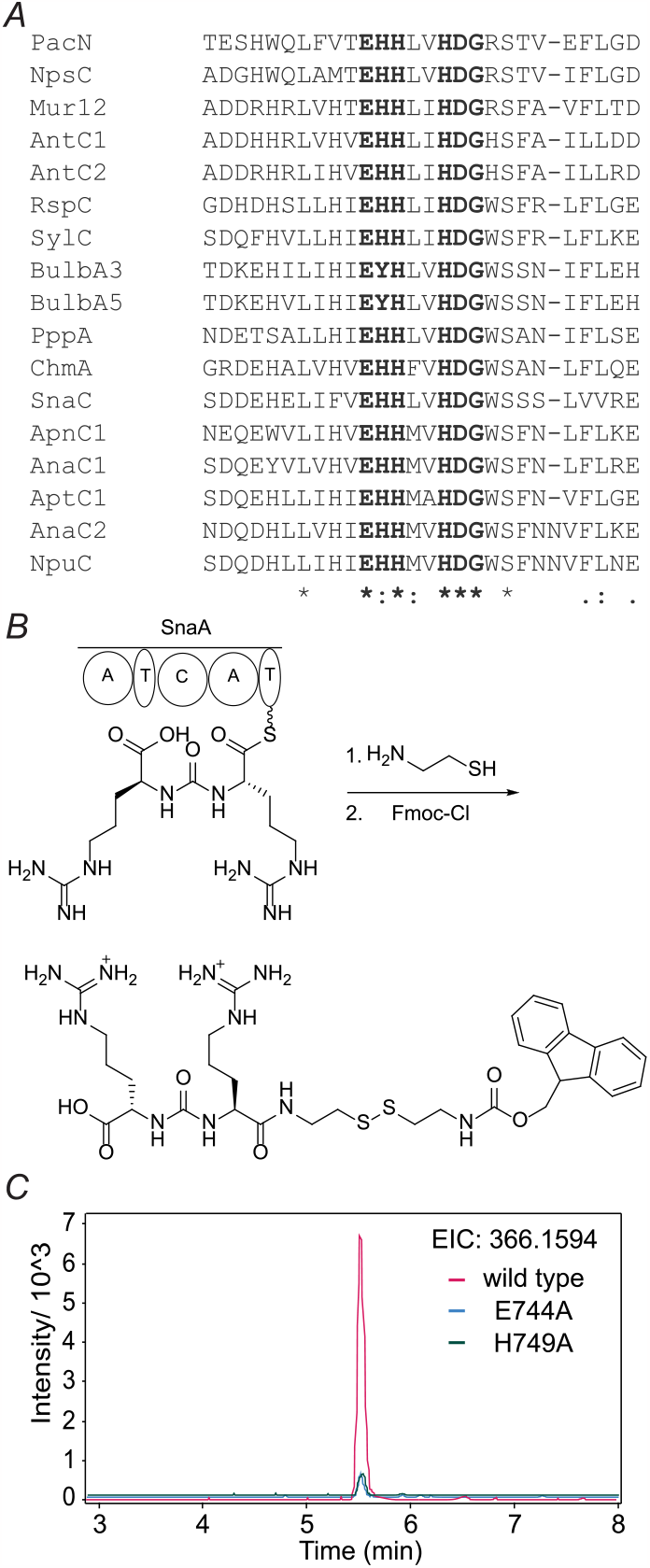
(*A*) Multiple sequence alignment of UreaC domains. The *Supporting Information* Table S3 contains the MiBiG BGC repository identification numbers for the listed enzymes. (*B*) Scheme of the derivatization of the bisarginine ureido structure generated by SnaA in vitro. (*C*) EICs of Fmoc- and cysteamine-derivatized bisarginine ureido structures generated in vitro by wild type and mutant SnaA.

The ureido-forming activity of SnaA was confirmed in vitro using holo-SnaA hetereologously expressed in *E. coli* BAP1.^43^ The PCP-bound products of SnaA were intercepted using cysteamine^44^ followed by chemical derivatization with fluorenylmethyloxycarbonyl chloride (Fmoc-Cl) and LC-MS analysis^45^ (Figure 4B). The observed products confirmed that SnaA is responsible for formation of intermediate **I** (Figure 3). When the active site glutamate and histidine residues of the UreaC domain were individually mutated to alanine, the resulting mutants showed significantly decreased production of the arginine ureido dipeptide in vitro (Figure 4C), indicating that the conserved glutamate and histidine residues in the UreaC active site are important (but not essential) for the ureido bond-forming activity of SnaA.

## Discussion

The formation of a ureido group during NRP biosynthesis is termed a chain-reversal event because it generates a carboxylate rather than the usual amine group at the initiation position. Proposed mechanisms of ureido group formation are presented in Figure S6. For ureido-forming BGCs that contain two A domains, the specificity of the A domain of the loading A-T didomain usually corresponds to the amino acid at the terminal position of the ureido group.^37,38,40,42^ Similarly, the A domain specificity of the first extension module (UreaC-A-T) usually corresponds to the internal amino acid of the ureido group.^37,38,40,42^ After ureido bond formation, the amino acid at the internal position is still attached to the PCP as a thioester and can be further extended by NRPS/PKS biochemistry.^42^ However, the terminal amino acid is left with an unactivated carboxylate and can no longer be extended by the assembly line chemistry.^42^ This model explains why all isolated ureido-containing NRPs only have one side of the ureido moiety further extended by the NRPS/PKS (Figure S7). If chain extension of the terminal carboxylate is desired for biological activity, two possible solutions can be envisioned. Either the ureido forming process will need to take place using a T-domain bound polypeptide (rather than amino acid) that is activated by ATP (Figure S6C), a mechanism that has been ruled out in the case of SylC.^27^ Alternatively, the system needs a separate amidation machinery. The PEARL-like enzyme SnaE appears to have been recruited for this latter purpose. Based on its sequence homology to PEARLs, SnaE is likely to add threonine using a similar ATP- and aminoacyl-tRNA-dependent mechanism,^17^ in which ATP is used to phosphorylate the terminal carboxylate to form an activated acyl-phosphate intermediate, which is then attacked by Thr-tRNA^Thr^ as the Thr donor in a condensation reaction. Hydrolysis of the tRNA as is observed in PEARLs would then provide the observed products.

The formation of the ethyl ketone in compound D and E follows a unique mechanism where the ethyl group originates from the decarboxylation of methylmalonate. Ethyl ketones are commonly observed motifs during PK/NRP biosynthesis, but the biosynthetic precursors of the ethyl group are usually *S*-adenosyl methionine (SAM) and malonate. For instance, the ethyl ketone derivative of arginine is a biosynthetic intermediate of saxitoxin^32,33^ and is biosynthesized by a polyketide-like synthase SxtA.^46^ Malonyl-CoA is loaded onto the ACP and is then methylated by the methyltransferase domain of SxtA. The methylmalonyl-ACP is thought to be decarboxylated to propionyl-ACP, which is followed by a pyridoxal phosphate-dependent condensation between arginine and propionyl-ACP to yield the arginine ethyl ketone.^46^

In the case of epoxyketone proteasome inhibitors such as epoxomicin and eponemycin,^47-49^ the ethyl groups of the epoxyketone warhead originate similarly from malonyl-CoA and on-ACP methylation(s) by a methyltransferase domain of the PKS EpxE/EpnH.^35,50^ The epoxide is thought to be generated by a conserved acyl-CoA dehydrogenase-like enzyme EpxF/EpnF via a decarboxylation-dehydrogenation-epoxidation sequence after thioesterase-mediated release from the assembly lines.^35^ Given that the vinylketone in compounds C/F originates from methylmalonate, the acyl-CoA dehydrogenase-like enzyme SnaO may also use a decarboxylative dehydrogenation mechanism to install the α,β-unsaturated ketone. Interestingly, the reaction of SnaO seems to stop at dehydrogenation, because no epoxidation was observed during heterologous expression in *S. albus*.

Different from the biosynthetic pathways of saxitoxin and epoxyketones, the methyltransferase domain for the methylation of malonate is absent in the *sna* BGC. This absence is consistent with the PKS SnaB using methylmalonyl-CoA to produce the ethyl group of threopeptin. The ethyl group of the epoxyketone macyranone could also originate from methylmalonate since its biosynthetic PKS module lacks a methyltransferase domain.^48^

We hypothesize that compound F (threopeptin) is the final product of the *sna* BGC because its biosynthesis depends on SnaA, SnaO, and SnaE, and it carries an α,β-unsaturated ketone that could function as an electrophilic warhead. The antipain group of protease inhibitors^51^ structurally resembles threopeptin. The aldehyde of antipain covalently targets protease active site serine or cysteine residues and the vinyl ketone of threopeptin may similarly target a protease active site serine or cysteine residue via 1,4-conjugate addition. Although the instability of threopeptin prevented isolation and bioactivity testing, *S. nassauensis* may produce the compound to inhibit proteases of competitor or predator organisms after secretion by SnaT_1_ and T_2_.

## Data Availability Statement

The authors declare that the data supporting the findings of this study are available within the paper and its Supporting Information files, and at Mendeley Data, V1, doi: 10.17632/rjytc5c3cr.1 as well as from the corresponding author upon reasonable request.

## Supporting Information

Supporting Figures S1-S7, Supporting Tables S1-S3, NMR spectra of all new compounds, and accession numbers of all proteins discussed in this study.

## Notes

The authors declare no competing financial interest(s).

## Acknowledgments

We thank Dr. Lingyang Zhu in the School of Chemical Sciences NMR Lab for assistance in data interpretation and Professor Gregory Challis (University of Warwick) for helpful discussions.

## References

(1) van Wageningen, A. M.; Kirkpatrick, P. N.; Williams, D. H.; Harris, B. R.; Kershaw, J. K.; Lennard, N. J.; Jones, M.; Jones, S. J.; Solenberg, P. J. Sequencing and analysis of genes involved in the biosynthesis of a vancomycin group antibiotic. Chem. Biol. 1998, 5, 155–162.

(2) Shen, B.; Du, L.; Sanchez, C.; Edwards, D. J.; Chen, M.; Murrell, J. M. Cloning and characterization of the bleomycin biosynthetic gene cluster from Streptomyces verticillus ATCC15003. J. Nat. Prod. 2002, 65, 422–431.

(3) Koumoutsi, A.; Chen, X. H.; Henne, A.; Liesegang, H.; Hitzeroth, G.; Franke, P.; Vater, J.; Borriss, R. Structural and functional characterization of gene clusters directing nonribosomal synthesis of bioactive cyclic lipopeptides in Bacillus amyloliquefaciens strain FZB42. J. Bacteriol. 2004, 186, 1084–1096.

(4) Bushley, K. E.; Raja, R.; Jaiswal, P.; Cumbie, J. S.; Nonogaki, M.; Boyd, A. E.; Owensby, C. A.; Knaus, B. J.; Elser, J.; Miller, D.; Di, Y.; McPhail, K. L.; Spatafora, J. W. The genome of Tolypocladium inflatum: evolution, organization, and expression of the cyclosporin biosynthetic gene cluster. PLoS Genet. 2013, 9, e1003496.

(5) Sieber, S. A.; Marahiel, M. A. Molecular mechanisms underlying nonribosomal peptide synthesis: approaches to new antibiotics. Chem. Rev. 2005, 105, 715–738.

(6) Fischbach, M. A.; Walsh, C. T. Assembly-line enzymology for polyketide and nonribosomal peptide antibiotics: logic, machinery, and mechanisms. Chem. Rev. 2006, 106, 3468–3496.

(7) Reimer, J. M.; Eivaskhani, M.; Harb, I.; Guarné, A.; Weigt, M.; Schmeing, T. M. Structures of a dimodular nonribosomal peptide synthetase reveal conformational flexibility. Science 2019, 366, eaaw4388.

(8) Flugel, R. S.; Hwangbo, Y.; Lambalot, R. H.; Cronan, J. E., Jr.; Walsh, C. T. Holo-(acyl carrier protein) synthase and phosphopantetheinyl transfer in Escherichia coli. J. Biol. Chem. 2000, 275, 959–968.

(9) Mahariel, M. A.; Stachelhaus, T.; Mootz, H. D. Modular peptide synthetases involved in nonribosomal peptide synthesis. Chem. Rev. 1997, 2651-2674.

(10) Ortega, M. A.; Hao, Y.; Zhang, Q.; Walker, M. C.; van der Donk, W. A.; Nair, S. K. Structure and mechanism of the tRNA-dependent lantibiotic dehydratase NisB. Nature 2015, 517, 509--512.

(11) Repka, L. M.; Chekan, J. R.; Nair, S. K.; van der Donk, W. A. Mechanistic understanding of lanthipeptide biosynthetic enzymes. Chem. Rev. 2017, 117, 5457–5520.

(12) Wang, S.; Wu, K.; Tang, Y. J.; Deng, H. Dehydroamino acid residues in bioactive natural products. Nat. Prod. Rep. 2023, doi: 10.1039/d1033np00041a.

(13) Montalbán-López, M.; Scott, T. A.; Ramesh, S.; Rahman, I. R.; van Heel, A. J.; Viel, J. H.; Bandarian, V.; Dittmann, E.; Genilloud, O.; Goto, Y.; Grande Burgos, M. J.; Hill, C.; Kim, S.; Koehnke, J.; Latham, J. A.; Link, A. J.; Martinez, B.; Nair, S. K.; Nicolet, Y.; Rebuffat, S.; Sahl, H.-G.; Sareen, D.; Schmidt, E. W.; Schmitt, L.; Severinov, K.; Süssmuth, R. D.; Truman, A. W.; Wang, H.; Weng, J.-K.; van Wezel, G. P.; Zhang, Q.; Zhong, J.; Piel, J.; Mitchell, D. A.; Kuipers, O. P.; van der Donk, W. A. New developments in RiPP discovery, enzymology and engineering. Nat. Prod. Rep. 2021, 138, 130 – 239.

(14) Hudson, G. A.; Zhang, Z.; Tietz, J. I.; Mitchell, D. A.; van der Donk, W. A. In vitro biosynthesis of the core scaffold of the thiopeptide thiomuracin. J. Am. Chem. Soc. 2015, 137, 16012–16015.

(15) Garg, N.; Salazar-Ocampo, L. M.; van der Donk, W. A. In vitro activity of the nisin dehydratase NisB. Proc. Natl. Acad. Sci. U. S. A. 2013, 110, 7258–7263.

(16) Ting, C. P.; Funk, M. A.; Halaby, S. L.; Zhang, Z.; Gonen, T.; van der Donk, W. A. Use of a scaffold peptide in the biosynthesis of amino acid-derived natural products. Science 2019, 365, 280–284.

(17) Zhang, Z.; van der Donk, W. A. Nonribosomal peptide extension by a peptide amino-acyl tRNA ligase. J. Am. Chem. Soc. 2019, 141, 19625–19633.

(18) Daniels, P. N.; Lee, H.; Splain, R. A.; Ting, C. P.; Zhu, L.; Zhao, X.; Moore, B. S.; van der Donk, W. A. A biosynthetic pathway to aromatic amines that uses glycyl-tRNA as nitrogen donor. Nat. Chem. 2022, 14, 71–77.

(19) Miyanaga, A.; Janso, J. E.; McDonald, L.; He, M.; Liu, H.; Barbieri, L.; Eustaquio, A. S.; Fielding, E. N.; Carter, G. T.; Jensen, P. R.; Feng, X.; Leighton, M.; Koehn, F. E.; Moore, B. S. Discovery and assembly-line biosynthesis of the lymphostin pyrroloquinoline alkaloid family of mTOR inhibitors in Salinispora bacteria. J. Am. Chem. Soc. 2011, 133, 13311–13313.

(20) Jordan, P. A.; Moore, B. S. Biosynthetic pathway connects cryptic ribosomally synthesized posttranslationally modified peptide genes with pyrroloquinoline alkaloids. Cell Chem. Biol. 2016, 23, 1504–1514.

(21) Yu, Y.; van der Donk, W. A. Biosynthesis of 3-thia-α-amino acids on a carrier peptide. Proc. Natl. Acad. Sci. U. S. A. 2022, 119, e2205285119.

(22) Du, L.; Shen, B. Biosynthesis of hybrid peptide-polyketide natural products. Curr. Opin. Drug Discov. Devel. 2001, 4, 215–228.

(23) Singh, M.; Chaudhary, S.; Sareen, D. Non-ribosomal peptide synthetases: Identifying the cryptic gene clusters and decoding the natural product. J. Biosci. 2017, 42, 175–187.

(24) Jenke-Kodama, H.; Dittmann, E. Bioinformatic perspectives on NRPS/PKS megasynthases: advances and challenges. Nat. Prod. Rep. 2009, 26, 874–883.

(25) Dekimpe, S.; Masschelein, J. Beyond peptide bond formation: the versatile role of condensation domains in natural product biosynthesis. Nat. Prod. Rep. 2021, 38, 1910–1937.

(26) Bloudoff, K.; Schmeing, T. M. Structural and functional aspects of the nonribosomal peptide synthetase condensation domain superfamily: discovery, dissection and diversity. Biochim. Biophys. Acta - Proteins Proteom. 2017, 1865, 1587–1604.

(27) Imker, H. J.; Walsh, C. T.; Wuest, W. M. SylC catalyzes ureido-bond formation during biosynthesis of the proteasome inhibitor syringolin A. J. Am. Chem. Soc. 2009, 131, 18263–18265.

(28) Miller, B. R.; Gulick, A. M. Structural biology of nonribosomal peptide synthetases. Methods Mol. Biol. 2016, 1401, 3–29.

(29) Kautsar, S. A.; Blin, K.; Shaw, S.; Navarro-Muñoz, J. C.; Terlouw, B. R.; van der Hooti, J. J. J.; van Santen, J. A.; Tracanna, V.; Suarez Duran, H. G.; Pascal Andreu, V.; Selem-Mojica, N.; Alanjary, M.; Robinson, S. L.; Lund, G.; Epstein, S. C.; Sisto, A. C.; Charkoudian, L. K.; Collemare, J.; Linington, R. G.; Weber, T.; Medema, M. H. MIBiG 2.0: a repository for biosynthetic gene clusters of known function. Nucleic Acids Res. 2020, 48, D454–D458.

(30) Chater, K. F.; Wilde, L. C. Restriction of a bacteriophage of Streptomyces albus G involving endonuclease SalI. J. Bacteriol. 1976, 128, 644–650.

(31) Bai, C.; Zhang, Y.; Zhao, X.; Hu, Y.; Xiang, S.; Miao, J.; Lou, C.; Zhang, L. Exploiting a precise design of universal synthetic modular regulatory elements to unlock the microbial natural products in Streptomyces. Proc. Natl. Acad. Sci. U. S. A. 2015, 112, 12181–12186.

(32) Tsuchiya, S.; Cho, Y.; Konoki, K.; Nagasawa, K.; Oshima, Y.; Yotsu-Yamashita, M. Synthesis and identification of proposed biosynthetic intermediates of saxitoxin in the cyanobacterium Anabaena circinalis (TA04) and the dinoflagellate Alexandrium tamarense (Axat-2). Org. Biomol. Chem. 2014, 12, 3016–3020.

(33) Tsuchiya, S.; Cho, Y.; Yoshioka, R.; Konoki, K.; Nagasawa, K.; Oshima, Y.; Yotsu-Yamashita, M. Synthesis and identification of key biosynthetic intermediates for the formation of the tricyclic skeleton of saxitoxin. Angew. Chem. Int. Ed. 2017, 56, 5327–5331.

(34) Thottumkara, A. P.; Parsons, W. H.; Du Bois, J. Saxitoxin. Angew. Chem. Int. Ed. 2014, 53, 5760–5784.

(35) Zabala, D.; Cartwright, J. W.; Roberts, D. M.; Law, B. J.; Song, L.; Samborskyy, M.; Leadlay, P. F.; Micklefield, J.; Challis, G. L. A flavin-dependent decarboxylase-dehydrogenase-monooxygenase assembles the warhead of α,β-epoxyketone proteasome inhibitors. J. Am. Chem. Soc. 2016, 138, 4342–4345.

(36) Zettler, J.; Zubeil, F.; Kulik, A.; Grond, S.; Kaysser, L. Epoxomicin and eponemycin biosynthesis involves gem-dimethylation and an acyl-CoA dehydrogenase-like enzyme. ChemBioChem 2016, 17, 792–798.

(37) Rouhiainen, L.; Jokela, J.; Fewer, D. P.; Urmann, M.; Sivonen, K. Two alternative starter modules for the non-ribosomal biosynthesis of specific anabaenopeptin variants in Anabaena (Cyanobacteria). Chem. Biol. 2010, 17, 265–273.

(38) Zhong, W.; Deutsch, J. M.; Yi, D.; Abrahamse, N. H.; Mohanty, I.; Moore, S. G.; McShan, A. C.; Garg, N.; Agarwal, V. Discovery and biosynthesis of ureidopeptide natural products macrocyclized via indole N-acylation in marine Microbulbifer spp. Bacteria. ChemBioChem 2023, 24, e202300190.

(39) Liu, J.; Zhou, H.; Yang, Z.; Wang, X.; Chen, H.; Zhong, L.; Zheng, W.; Niu, W.; Wang, S.; Ren, X.; Zhong, G.; Wang, Y.; Ding, X.; Müller, R.; Zhang, Y.; Bian, X. Rational construction of genome-reduced Burkholderiales chassis facilitates efficient heterologous production of natural products from proteobacteria. Nat. Commun. 2021, 12, 4347.

(40) Ióca, L. P.; Dai, Y.; Kunakom, S.; Diaz-Espinosa, J.; Krunic, A.; Crnkovic, C. M.; Orjala, J.; Sanchez, L. M.; Ferreira, A. G.; Berlinck, R. G. S.; Eustáquio, A. S. A family of nonribosomal peptides modulate collective behavior in Pseudovibrio bacteria isolated from marine sponges. Angew. Chem. Int. Ed. 2021, 60, 15891– 15898.

(41) Zhang, W.; Ostash, B.; Walsh, C. T. Identification of the biosynthetic gene cluster for the pacidamycin group of peptidyl nucleoside antibiotics. Proc. Natl. Acad. Sci. U. S. A. 2010, 107, 16828–16833.

(42) Cui, Z.; Nguyen, H.; Bhardwaj, M.; Wang, X.; Büschleb, M.; Lemke, A.; Schütz, C.; Rohrbacher, C.; Junghanns, P.; Koppermann, S.; Ducho, C.; Thorson, J. S.; Van Lanen, S. G. Enzymatic C(β)-H functionalization of L-Arg and L-Leu in nonribosomally derived peptidyl natural products: a tale of two oxidoreductases. J. Am. Chem. Soc. 2021, 143, 19425–19437.

(43) Pfeifer, B. A.; Admiraal, S. J.; Gramajo, H.; Cane, D. E.; Khosla, C. Biosynthesis of complex polyketides in a metabolically engineered strain of E. coli. Science 2001, 291, 1790–1792.

(44) Belecki, K.; Townsend, C. A. Biochemical determination of enzyme-bound metabolites: preferential accumulation of a programmed octaketide on the enediyne polyketide synthase CalE8. J. Am. Chem. Soc. 2013, 135, 14339–14348.

(45) Patteson, J. B.; Dunn, Z. D.; Li, B. In vitro biosynthesis of the nonproteinogenic amino acid methoxyvinylglycine. Angew. Chem. Int. Ed. 2018, 57, 6780–6785.

(46) Chun, S. W.; Hinze, M. E.; Skiba, M. A.; Narayan, A. R. H. Chemistry of a unique polyketide-like synthase. J. Am. Chem. Soc. 2018, 140, 2430–2433.

(47) Schorn, M.; Zettler, J.; Noel, J. P.; Dorrestein, P. C.; Moore, B. S.; Kaysser, L. Genetic basis for the biosynthesis of the pharmaceutically important class of epoxyketone proteasome inhibitors. ACS Chem. Biol. 2014, 9, 301–309.

(48) Keller, L.; Plaza, A.; Dubiella, C.; Groll, M.; Kaiser, M.; Müller, R. Macyranones: structure, biosynthesis, and binding mode of an unprecedented epoxyketone that targets the 20S proteasome. J. Am. Chem. Soc. 2015, 137, 8121–8130.

(49) Owen, J. G.; Charlop-Powers, Z.; Smith, A. G.; Ternei, M. A.; Calle, P. Y.; Reddy, B. V.; Montiel, D.; Brady, S. F. Multiplexed metagenome mining using short DNA sequence tags facilitates targeted discovery of epoxyketone proteasome inhibitors. Proc. Natl. Acad. Sci. U S A 2015, 112, 4221–4226.

(50) Liu, J.; Zhu, X.; Zhang, W. Identifying the minimal enzymes required for biosynthesis of epoxyketone proteasome inhibitors. ChemBioChem 2015, 16, 2585–2589.

(51) Suda, H.; Aoyagi, T.; Hamada, M.; Takeuchi, T.; Umezawa, H. Antipain, a new protease inhibitor isolated from actinomycetes. J. Antibiot. 1972, 25, 263–266.

