## Supporting Information for "PEARL-catalyzed peptide bond formation after chain reversal during the biosynthesis of non-ribosomal peptides"

**General Methods:**

All oligonucleotides used in this study were purchased from Integrated DNA Technologies. Adenosine-5'-triphosphate, apramycin, kanamycin, and isopropyl β-d-1-thiogalactopyranoside were purchased from GoldBio. Sodium acetate-$^{13}C_2$, cysteamine, magnesium chloride, and fluorenylmethyloxycarbonyl chloride (Fmoc-Cl) were purchased from Sigma-Aldrich. Sodium propionate (2-$^{13}$C) was purchased from Cambridge Isotope Laboratories. Liquid chromatography-high-resolution mass spectrometry data was recorded on an Agilent 1260 Infinity II coupled to a diode array detector WR and a G6545B mass spectrometer. Purine and HP-0921 (Agilent Technologies, Santa Clara, CA, USA) were used as reference masses during acquisition. Spectra were processed using the Agilent MassHunter Workstation Qualitative Analysis software (version 10.0). Nuclear magnetic resonance data was either acquired on a Bruker Avance III HD equipped with a 500-MHz, 5-mm, BBFO CryoProbe or a spectrometer equipped with a Bruker Avance Neo console and 600-MHz, 5-mm, BBO-BB Prodigy CryoProbe. NMR spectra were referenced to residual NMR solvent signals of dimethylsulfoxide (2.50 ppm, $^{1}$H; 39.5 ppm, $^{13}$C).

**Cloning of *sna* BGC and the selective inactivation of biosynthetic enzymes:**

The 17-kilobase (kb) BGC was divided into three fragments for amplification from the genomic DNA of *Stackebrandtia nassauensis* by NEB Q5 high-fidelity DNA polymerase using the primer pairs Sna1F/R, Sna2F/R, and Sna3F/R (Table S1). The expression vector pOSV801 was linearized using the primer pairs Sna_OSVF/R. The plasmid was assembled from three BGC DNA fragments and a linear fragment of pOSV801 with a strong promoter SP44[1] using Gibson Assembly.[2] The sequence of the plasmid termed pOSV801-SP44-Sna was verified using Sanger sequencing by primer walking.

The DNA sequence of SP44:
tgttcacattcgaaccgtctctgctttgacaacatgctgtgcggtgttgtaaagtctggtgtgaccctaacgaggagatcggttcacccat

For inactivation of SnaA during heterologous expression, both active site serine codons (S556 and S1616) in the two peptidyl carrier proteins were mutated to alanine (GCC). pOSV801-SP44-Sna was digested with NheI (ThermoFisher FD0974) and FspAI (ThermoFisher FD1664), and the 15-kb digested fragment without SnaA was purified by DNA gel electrophoresis. The sequence of the other fragment containing SnaA was divided into three pieces based on the location of the conserved serine codon of the PCP. The three fragments were amplified by Q5 DNA polymerase, and alanine mutations were introduced by the primer pairs $SnaA_{in}$_1F/R, $SnaA_{in}$_2F/R, and

$SnaA_{in}$_3F/R (Table S1). The 15-kb fragment without SnaA from restriction digestion and the three PCR fragments with alanine mutations were assembled by Gibson assembly.

For inactivation of SnaB during heterologous expression, the active site serine (S927) in the acyl carrier protein (ACP) was mutated to alanine (GCC). pOSV801-SP44-Sna was digested with EcoRV (NEB R0195S) and FspAI (ThermoFisher FD1664), and the 14-kb digested fragment without the ACP of SnaB was purified by DNA gel electrophoresis. The rest of the plasmid was amplified using primer pairs $SnaB_{in}$_1F/R and $SnaB_{in}$_2F/R (Table S1). The primers similarly introduced the alanine mutation during PCR. The 14-kb fragment without the ACP of SnaA and the two fragments with alanine mutations were assembled by Gibson assembly.

For the inactivation of SnaE and SnaO, in-frame deletion was used. The open reading frame of SnaE was shortened from 2538 bp to 270 bp by retaining only the first 135 base pair (bp) and the last 135 bp of the DNA sequence of SnaE. This will result in a nonfunctional SnaE transcript in the expression plasmid. The expression plasmid with SnaE inactivation was constructed using a 3-fragment Gibson assembly after amplification of pOSV801-SP44-Sna using primer pairs $SnaE_{in}$_1F/R, $SnaE_{in}$_2F/R, and $SnaE_{in}$_3F/R (Table S1).

The inactivation of SnaO was similarly achieved by retaining the first 150 bp and the last 129 bp of its DNA sequence. This truncates the open reading frame of SnaO from 1662 bp to 279 bp and will result in a nonfunctional SnaO. The parent plasmid pOSV801-SP44-Sna was digested by EcoRV (NEB R0195S) and FspAI (ThermoFisher FD1664) and the 14-kb fragment without SnaO was purified by DNA gel electrophoresis. The expression plasmid with SnaO inactivation was constructed using a 3-fragment Gibson assembly from the 14-kb EcoRV-FspAI fragment and Two fragments from the amplification of pOSV801-SP44-Sna using primer pairs $SnaO_{in}$_1F/R, $SnaO_{in}$_2F/R (Table S1).

**Metabolomic identification of the products of *sna* BGC:**

The expression plasmid for the *sna* BGC and derivatives thereof were used to transform *E. coli* ET12567/pUZ8002. The plasmids were then transferred into *S. albus* J1074 by intergeneric conjugation according to previously described procedures.[3] In an 18×150 mm culture tube, 3 μL of spore stocks of *S. albus* exconjugants were inoculated into 4 mL of R5 medium.[4] The culture was grown on a rotating tube roller at 30 °C for 4 days. After 4 days, the supernatant and the cell pellet were separated by centrifugation under 4500 rpm for 5 min. The spent medium supernatant was purified using a 100 mg C18 solid phase extraction column (ThermoFisher 60108-302). The column was conditioned with methanol and equilibrated with 2 column volumes (CV) of $H_2O$. The medium was passed through the C18 SPE column using a vacuum manifold. The SPE column was again washed with 2 CV of $H_2O$. Bound metabolites were eluted using 350 μL of 50% acetonitrile in $H_2O$ and 150 μL of acetonitrile. The cell pellet was lyophilized to dryness and was extracted with 1 mL of acetone. The acetone extract was evaporated using a SpeedVac vacuum concentrator (SPD140DDA), and the residue was redissolved in 200 μL of MeOH.

The spent medium extract was analyzed on an Agilent 1260 Infinity II coupled with a G6545B liquid chromatography-mass spectrometry (LC-MS) instrument. Each sample was analyzed on a polar C18 column (Phenomenex 00F-4759-AN) maintained at 45 °C using gradient one and on a hydrophilic interaction chromatography (HILIC) column (Phenomenex 00B-4461-AN) maintained at 40 °C using gradient two.

Gradient one:

Solvent A: $H_2O$ + 0.1% formic acid. Solvent B: acetonitrile + 0.1% formic acid

0% B at 0-2 min; 0% B to 60% over 18 min; 60% B to 90% B over 4 min; 90% B to 95% B over 0.5 min; 95% B to 0% B over 1 min. The column was re-equilibrated at 0% B between injections for 4 min.

Gradient two:

Solvent A: $H_2O$ + 10 mM ammonium formate, pH 3.2. Solvent B: 95% acetonitrile in $H_2O$ + 10 mM ammonium formate, pH 3.2.

100% B at 0-1 min; 100% B to 60% over 12 min; 60% B to 100% B over 0.5 min. The column was re-equilibrated at 100% B between injections for 2.8 min.

MS settings:

ion polarity: positive; mass range: 50-1700 m/z; slicer mode: high resolution, 2 GHz; gas temperature: 325 °C; drying gas: 10 L/min; nebulizer pressure: 35 psi; sheath gas temperature 375 °C; sheath gas flow 11 L/ min; capillary voltage: 3500 V; nozzle voltage: 0 V; fragmentor voltage: 120 V; skimmer: 65 V; Oct 1 RF Vpp: 750 V. Acquisition mode: MS. Acquisition rate: 5 Hz.

Three biological replicates of the empty plasmid pOSV801 exconjugants and the *sna* BGC exconjugants were analyzed in parallel. The Agilent data files were converted to mzML data formats using MSConvert.[5] Comparative metabolomics calculations were performed using XCMS[6] pairwise comparison with UPLC/UHD Q-TOF default parameters. New metabolites were predominantly in the medium supernatant. The metabolites were identified using either C18 or HILIC columns. However, the HILIC column offered better MS response and separation of the target metabolites from medium components and, therefore, was chosen for analysis.

**Isolation of compounds A, B, and C for structural elucidation:**

Eight microliters of spores of *S. albus sna* BGC exconjugants were inoculated into 50 mL of TSB medium in 250 mL flasks with stainless steel springs as baffles. This starter culture was incubated at 30 °C 220 rpm for 2 days. The starter culture was inoculated into 1 L of R5 medium in a 4 L baffled flask and was incubated at 30 °C and 220 rpm for another 4 days. The spent medium was harvested by centrifugation at 5000 g for 15 min and was further clarified by filtration against filter paper. A 10 g C18 SPE column (ThermoFisher 60108-703) hydrated with MeOH and equilibrated with $H_2O$ was used to extract 150 mL of the spent medium. After washing with 2 CV of $H_2O$, bound metabolites were eluted using 2×10 mL 25% acetonitrile in $H_2O$, 2×10 mL 50% acetonitrile in $H_2O$, and 2×10 mL acetonitrile. The elution was concentrated by lyophilization and was further purified using a 10 mm × 250 mm Atlantis HILIC column with the following gradient:

Solvent A: $H_2O$ + 10 mM ammonium formate, pH 3.2. Solvent B: 95% acetonitrile in $H_2O$ + 10 mM ammonium formate, pH 3.2. Flow rate: 4 mL/ min.

100% B to 80% over 8 min; 80% B to 64% B over 14 min, 64% B to 60% B over 2 min, 60% B to 100% B over 1 min. The column was re-equilibrated at 100% B between injections for 5 min.

Metabolites from the *sna* BGC eluted from 18-24 min. The elutions were combined and lyophilized to dryness. The residual solid after lyophilization was resuspended in $H_2O$ and was further purified on a 4.6 × 250 mm Vydac C18 column using the following gradient:

Solvent A: $H_2O$ + 0.1% trifluoroacetic acid. Solvent B: acetonitrile + 0.1% trifluoroacetic acid

2% B to 20% over 18 min; 20% B to 60% B over 2 min; 60% B to 2% B over 1 min. The column was re-equilibrated at 2% B between injections for 4 min.

Compound A eluted at 12.4 min. Compound B eluted at 12.8 min. Compound C eluted at 13.2 min. The elution fraction for each compound was lyophilized to dryness for further characterization.

#### $^{13}$C labeling of the product of polyketide synthase SnaB

Stock solutions of 100 mg/mL $^{13}C_2$-sodium acetate and 2-$^{13}$C-sodium propionate were prepared in $H_2O$ and were sterile-filtered. To an 18×150 mm culture tube were added 4 mL of R5 medium and 40 μL of 100 mg/mL $^{13}C_2$-sodium acetate or 2-$^{13}$C-sodium propionate. Three microliters of *S. albus* exconjugants spore suspension were inoculated into the expression medium. The culture supernatant was harvested after four days and purified as described above using a 100 mg C18 SPE column.

#### Sequence analysis of ureido-generating condensation domains

The BGCs of ureido-containing NRPs were compiled from MiBiG or NCBI, including antipain (BGC0001570,[7] BGC0002051[8-9]), anabaenopeptins (BGC0000301,[10] BGC0000302,[11] BGC0001479,[12-13] BGC0002512[14]), syringolin (BGC0000436,[15] BGC0001047[16]), pacidamycin (BGC0000951[17]), napsamycin (BGC0000950[18-19]), muraymycin (BGC0001020[9, 20]), chitinimide (BGC0002503[21]), pseudovibriamide (BGC0002123[22]), and bulbiferamide (NCBI Bioproject PRJNA941849).[23-24] The C domain sequences were extracted based on the domain annotation in MiBiG.

By matching the adenylation domain specificity and the amino acids on each side of the ureido moiety of the final product, the UreaC domain were identified based on the canonical C-A-T module architecture. In modular NRPSs, the UreaC domain was always found in the first peptide extension module following the loading A-T didomain, as in the case of anabaenopeptins, chitinimide, pseudovibriamide, and bulbiferiamide.

The UreaC domains in syringolin A,[25] pacidamycin,[17] antipain,[9] and muraymycin[9] have been confirmed by in vitro enzyme activity, which allowed the prediction of UreaC domains from similar BGCs such as napsamycin[18] and deimino-antipain.[7]

The amino acid translations of all genes in MiBiG were downloaded in a FASTA file (Version 3.1). The condensation domain sequences were filtered out by searching the database using the hidden Markov model of PF00668 with an E value threshold of 0.01. Examining all the extracted condensation domain sequences showed that the EHHXXHDG active site is confined to UreaC domains.

#### Reconstitution of SnaA enzymatic activity in vitro.

The DNA sequence of SnaC was amplified from pOSV801-SP44-Sna using the primer pair His-TEV-SnaC_F and His-TEV-SnaC_R (Table S1). The expression vector pRSFDuet was linearized by the primer pair SnaC_RSF_F/R. SnaC was cloned into the multiple cloning site II of pRSFDuet by Gibson assembly to yield the plasmid construct: pRSFDuet: empty: $His_6$-TEV-SnaC.

The DNA sequence encoding SnaA was codon-optimized and synthesized by Twist Biosciences in three fragments. The three synthetic gene fragments of *snaA* were amplified using the primer pairs coSnaA1_F/R, coSnaA2_F/R, and coSnaA3_F/R (Table S1). The expression vector pRSFDuet: empty: $His_6$-TEV-SnaC was linearized by the primer pair coSnaA_RSF_F/R. The DNA fragments of SnaA were inserted into the multiple cloning site I of pRSFDuet: empty: $His_6$-TEV-SnaC with a C-terminal Histag using Gibson assembly to yield the plasmid pRSFDuet: coSnaA-TEV-$His_6$: $His_6$-TEV-SnaC. The UreaC domain active site mutants E744A and H749A were generated using the NEB KLD method with the primer pairs coSnaAE744A_F/R and coSnaAH749A_F/R (Table S1).

The above plasmid containing SnaA and SnaC was used to transform *E. coli* BAP1 cells for protein expression. The transformants were selected on LB agar with 50 μg/mL kanamycin at 37 °C. A single colony was used to inoculate 7 mL of LB with 50 μg/mL kanamycin and was grown at 37 °C and 220 rpm for 4 h. The seed culture was used to inoculate 500 mL of LB with 50 μg/mL kanamycin, 1 mM $MgSO_4$, and trace metal mix (1X, Teknova T1001). The culture was grown at 37 °C and 220 rpm until $OD_{600}$= 0.6 when the shaker temperature was reduced to 16 °C. After 30 min of incubation at 16 °C, protein expression was induced by the addition of 0.2 mM final concentration of IPTG.

The cells were harvested after 16 h of incubation by centrifugation under 5000 g at 10 °C for 15 min and were resuspended in protein purification buffer containing 25 mM 3-(N-morpholino)propanesulfonic acid (MOPS), 150 mM KCl, 10% glycerol, pH 7.5. The cells were lysed by sonication under 40% maximal amplitude with 1 s on and 2 s off for 3 min. The lysate was clarified by centrifugation at 49990 g at 10 °C for 15 min. The imidazole concentration of the supernatant was adjusted to 10 mM, and the supernatant was incubated with His60 Ni Superflow Resin (Takara 635660) at 4 °C for 20 min. The resin was harvested by centrifugation under 2000 g for 3 min and was resuspended in the protein purification buffer supplied with 30 mM imidazole. The resin suspension was transferred to a gravity-flow column, and the resin was further washed with 20 mL of protein purification buffer supplied with 30 mM imidazole and 5 mL of protein purification buffer supplied with 50 mM imidazole. Bound protein was eluted with 0.5 mL of 25 mM MOPS, 300 mM NaCl, 300 mM imidazole, 10% glycerol, pH 7.5 six times. The pooled elution fractions were concentrated and the buffer was exchanged to the protein purification buffer using an Amicon Ultra-4 Centrifugal Filter with 30 kDa molecular weight cutoff (MWCO). The protein solution containing SnaA and SnaC was flash-frozen with liquid nitrogen and was stored at -80 °C.

Ureido-peptide formation assay was performed in 50 μL of 50 mM HEPES, 100 mM NaCl (pH 7.5) containing 5 mM $NaHCO_3$, 4 mM $MgCl_2$, 0.5 mM tris(2-carboxyethyl)phosphine (TCEP), 1 mM arginine, 4 mM ATP, 5 μM of SnaA-TEV-$His_6$ coexpressed with $His_6$-TEV-SnaC, and 40 mM cysteamine (final concentration). Two hours later, 25 μL of the reaction solution was taken out and quenched by 50 μL cold acetonitrile. The remaining 25 μL of the reaction solution was quenched by 50 μL of cold acetonitrile 7 h after the setup. The quenched reaction was centrifuged at 20000 g for 3 min to remove the protein precipitates.

A 25 μL sample of the quenched reaction supernatant was mixed with 12.5 μL of 200 mM sodium borate buffer (pH 10.4) and 12.5 μL of 15 mg/mL fluorenylmethyloxycarbonyl chloride (Fmoc-Cl) in acetonitrile. The reaction was allowed to proceed at room temperature for 30 min. The derivatization reaction was centrifuged at 20000 g for 3 min to remove the precipitates, and 1 μL of the reaction supernatant was used for LC-MS analysis with the following condition.

Column: Agilent 3×100 mm Poroshell C18 maintained at 45 °C

Solvent A: $H_2O$ + 0.1% formic acid. Solvent B: acetonitrile + 0.1% formic acid

The gradient was 15% to 85% solvent B over 8 min. The column was re-equilibrated at 15% B between injections for 3 min.

NMR data of compound A (for actual spectra vide infra):

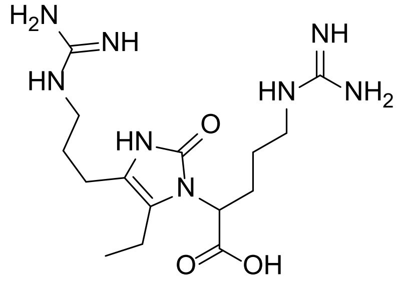

1H NMR (600 MHz, DMSO) δ 9.80 (s, 1H), 7.72 (br, 1H), 7.58 (br, 1H), 4.41 (t, *J* = 7.6 Hz, 1H), 3.14 – 3.04 (m, 4H), 2.36 – 2.22 (m, 4H), 2.04 (q, *J* = 8.3 Hz, 2H), 1.65 (p, *J* = 7.4 Hz, 2H), 1.47 – 1.41 (m, 1H), 1.32 – 1.26 (m, 1H), 0.97 (t, *J* = 7.4 Hz, 3H).

$^{13}C$ NMR (151 MHz, DMSO) δ 171.9, 156.7, 153.1, 119.8, 114.6, 53.6, 40.4, 40.2, 27.9, 26.3, 25.7, 20.7, 15.6, 14.8.

NMR data of compound B (for actual spectra vide infra):

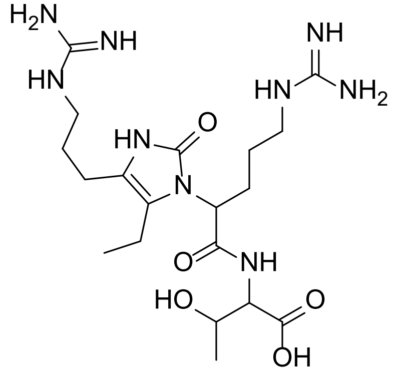

$^1H$ NMR (500 MHz, DMSO) δ 10.01 (s, 1H), 7.73 – 7.70 (m, 2H), 7.59 (t, *J* = 5.8 Hz, 1H), 5.01 (s, 1H), 4.38 (t, *J* = 8.0 Hz, 1H), 4.17 – 4.13 (m, 2H), 3.13 – 3.05 (m, 4H), 2.38 – 2.27 (m, 4H), 2.14 – 2.06 (m, 2H), 1.65 (p, *J* = 7.7 Hz, 2H), 1.47 – 1.38 (m, 1H), 1.33 – 1.27 (m, 1H), 1.03 (d, *J* = 6.2 Hz, 3H), 0.98 (t, *J* = 7.4 Hz, 3H).

$^{13}C$ NMR (126 MHz, DMSO) δ 171.8, 170.5, 156.8, 156.7, 153.6, 119.9, 115.5, 66.0, 57.6, 56.5, 40.3, 40.1, 27.8, 26.2, 25.7, 20.7, 20.4, 15.7, 14.7.

NMR data of compound C (for actual spectra vide infra):

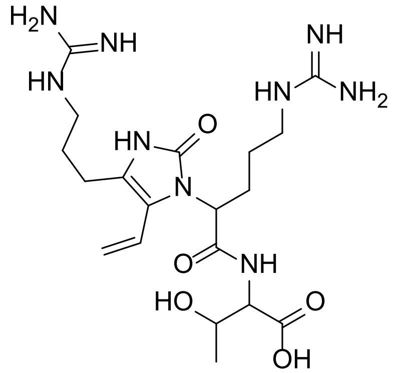


[1]H NMR (600 MHz, DMSO) δ 10.45 (s, 1H), 7.64 (br, 1H), 7.60 (d, J = 8.5 Hz, 1H), 7.56 (br, 1H), 6.37 (dd, J = 17.7, 11.6 Hz, 1H), 5.15 (dd, 2H), 4.99 (d, J = 6.6 Hz, 1H), 4.78 (dd, J = 10.8, 5.1 Hz, 1H), 4.18 (dd, J = 8.5, 3.1 Hz, 1H), 4.14 (br, 1H), 3.13 – 3.04 (m, 4H), 2.47 – 2.41 (m, 2H), 2.10 – 2.06 (m, 1H), 1.89 (br, 1H), 1.70 (p, J = 7.2 Hz, 2H), 1.35 – 1.21 (m, 2H), 1.01 (d, 3H).

[13]C NMR (151 MHz, DMSO) δ 171.8, 170.5, 156.6, 153.4, 124.1, 120.1, 117.5, 114.4, 66.1, 58.0, 55.0, 40.1, 40.1, 27.7, 26.6, 25.4, 21.5, 20.5.

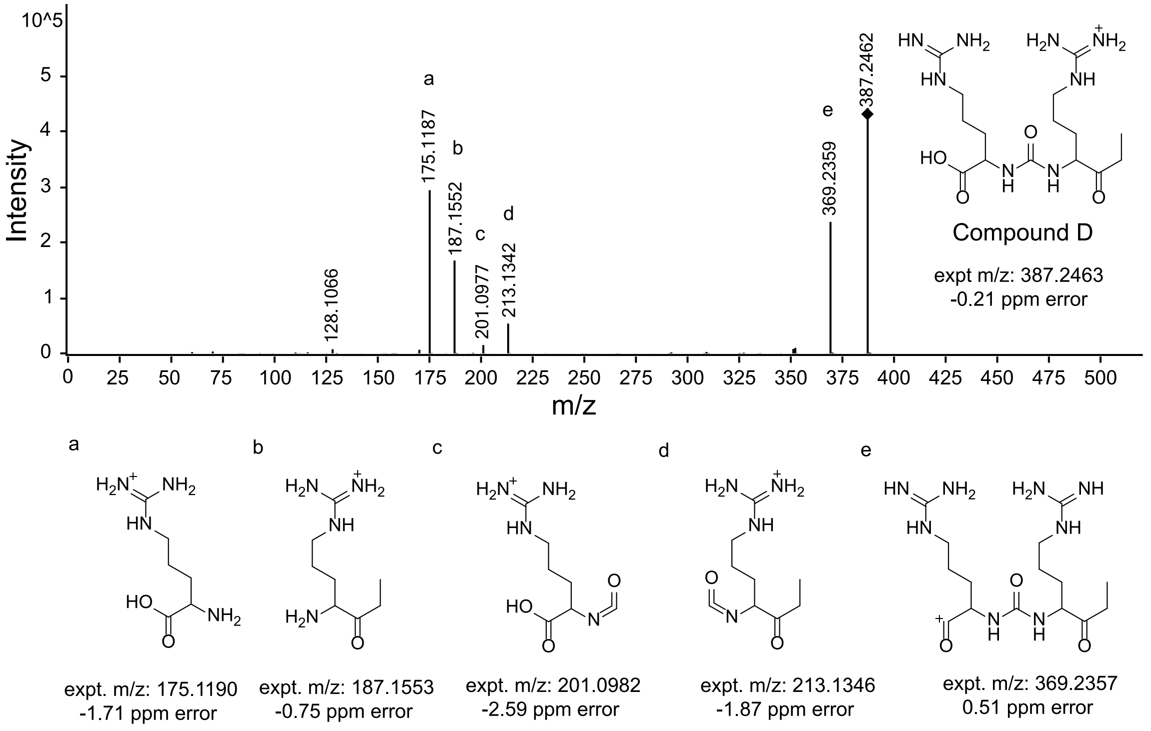


**Figure S1.** High-resolution tandem mass spectrometry (MS/MS) of compound D using 10 eV collision energy. The assignment of selected fragment ions and their ppm errors compared to calculated m/z values are listed.

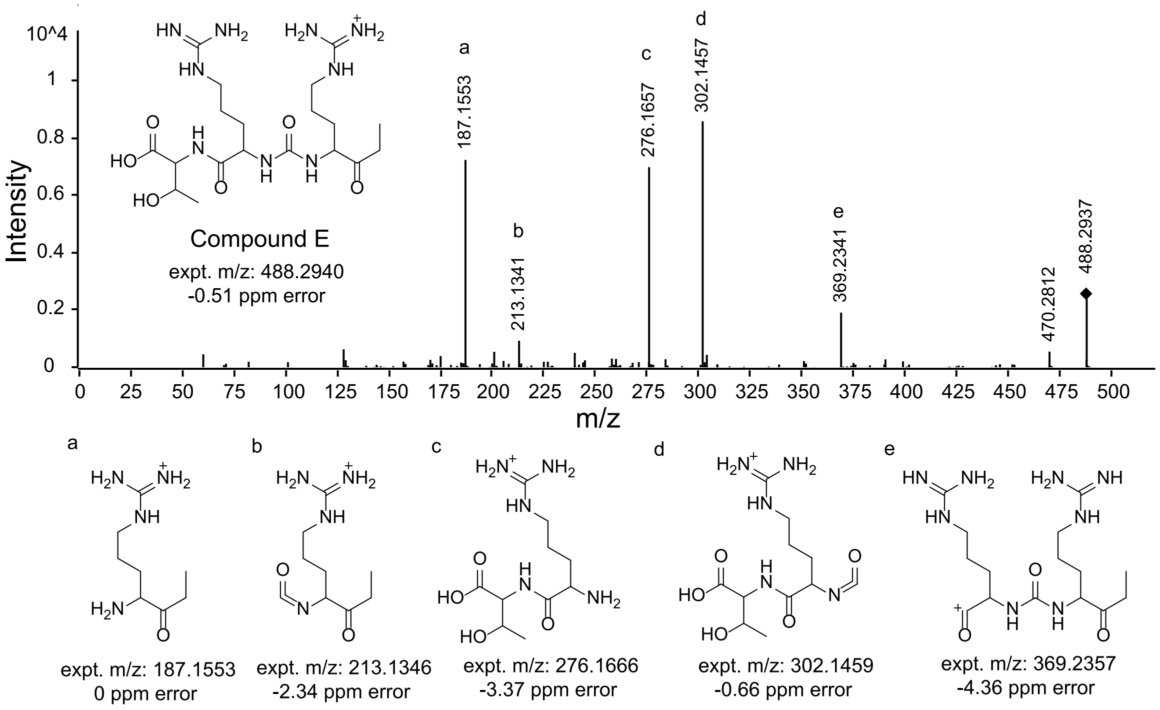


**Figure S2**. High-resolution tandem mass spectrometry (MS/MS) spectrum of compound E using 20 eV collision energy. The assignment of selected fragment ions and their ppm errors compared to calculated m/z values are listed.

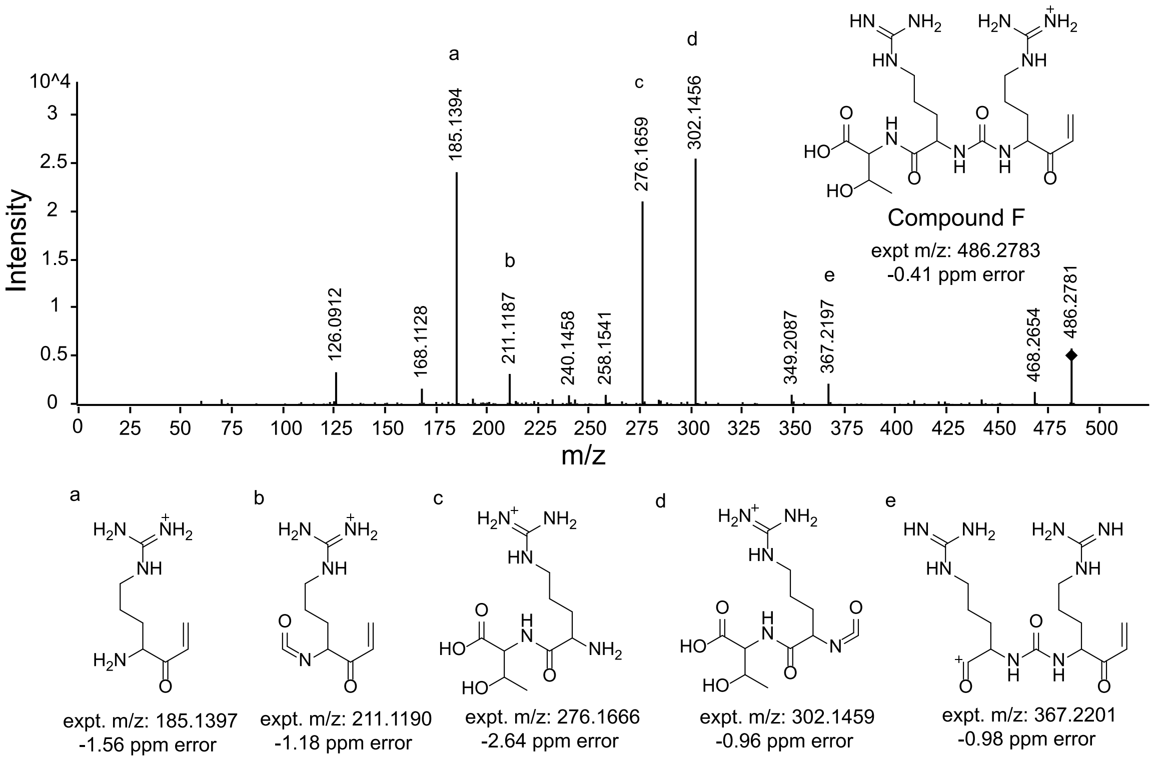


**Figure S3.** High-resolution tandem mass spectrometry (MS/MS) spectrum of compound F using 20 eV collision energy. The assignment of selected fragment ions and their ppm errors compared to calculated m/z values are listed.

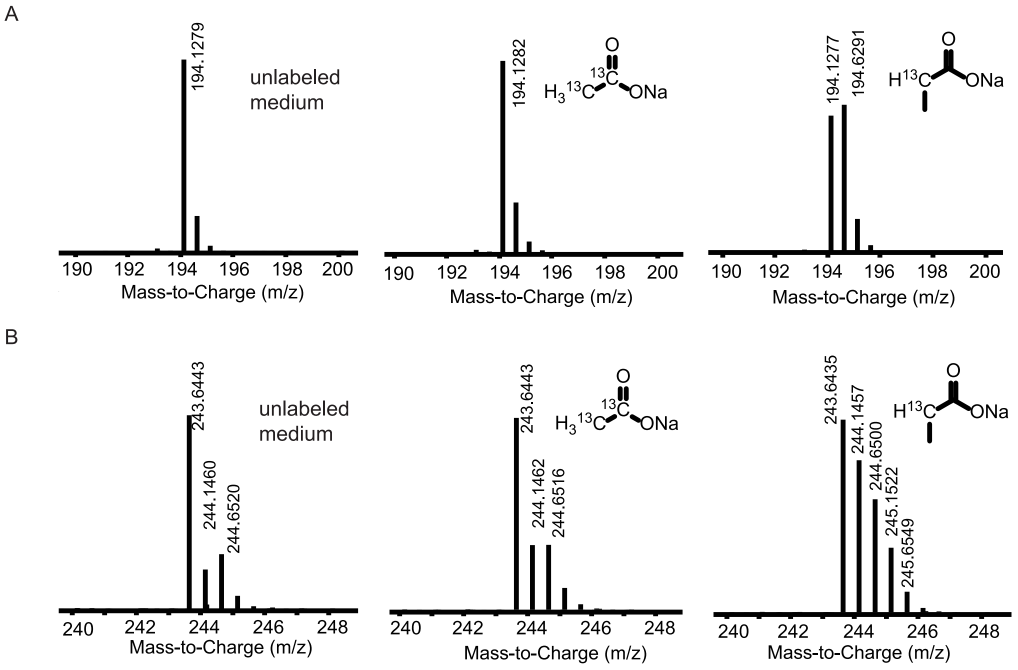


**Figure S4.** Comparison of MS spectra of compounds D, E, and F when different PKS precursors were fed. (*A*) MS spectra of compound D when the medium was supplied with $^{13}C_2$-sodium acetate or 2-$^{13}$C-sodium propionate. Only significant isotope incorporation was observed when the medium was supplied with 2-$^{13}$C-sodium propionate. These data are consistent with conversion of proprionate to methylmalonate and methylmalonyl-CoA, and subsequent utilization by SnaB. (*B*) Mass spectra of compounds E and F when the medium was supplied with $^{13}C_2$-sodium acetate or 2-$^{13}$C-sodium propionate. Only significant isotope incorporation was observed when the medium was supplied with 2-$^{13}$C-sodium propionate. Compound E (244.6506 m/z, [M+2H]) and F (243.6428 m/z, [M+2H]) coelute under the HILIC-MS condition, which explains the unusual isotope distribution of the spectra.

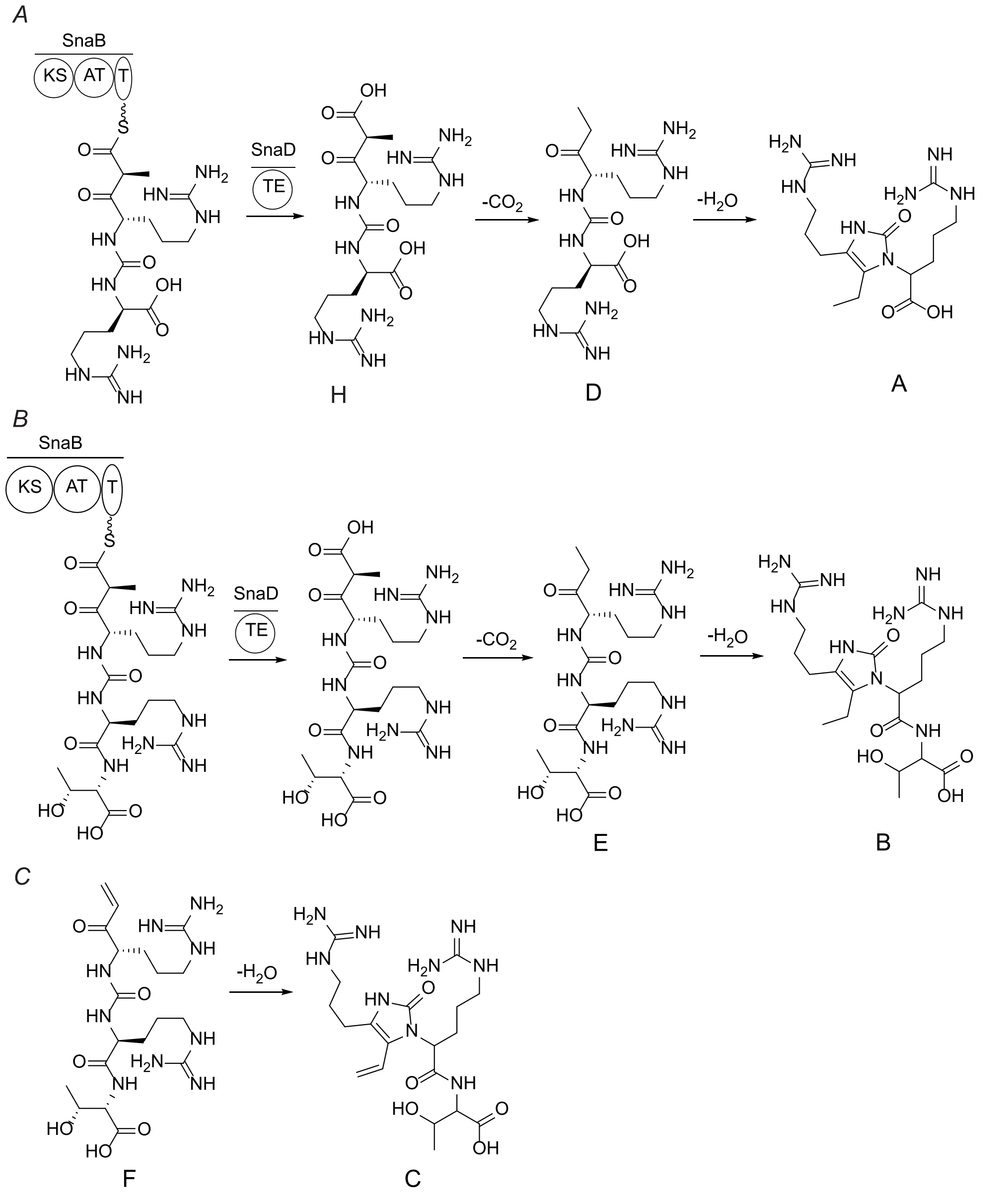


**Figure S5.** Proposed pathways towards the formation of compounds A-E. (*A*) Proposed pathway towards the formation of A and D. (*B*) Proposed pathway towards the formation of B and E. (*C*) Proposed pathway towards the formation of C. The pathways in panels B and C assume that the Thr is added prior to TE-catalyzed release from SnaB.

A.

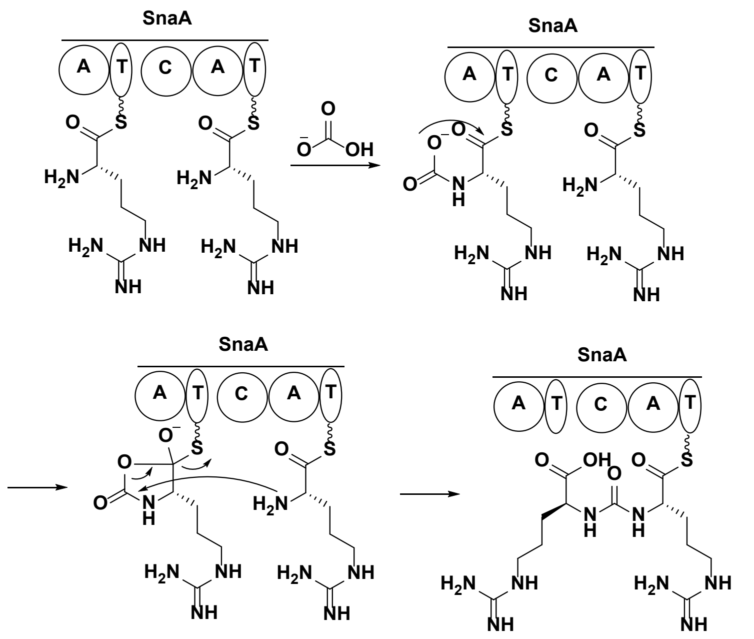


B.

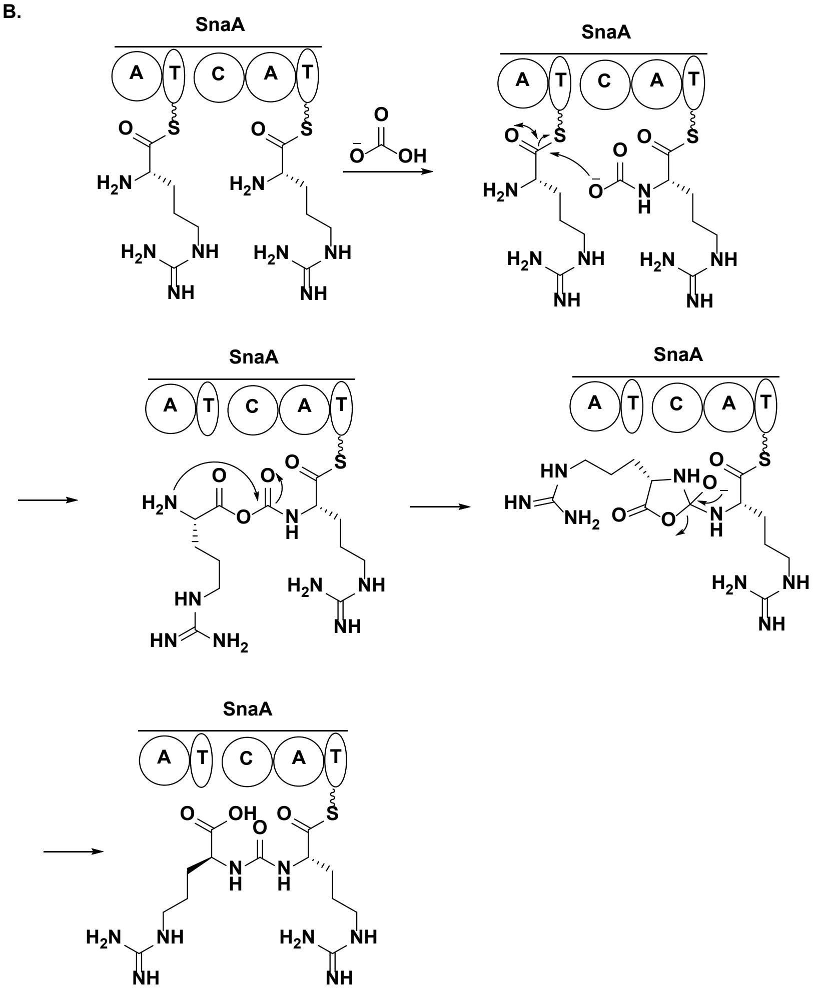

C.

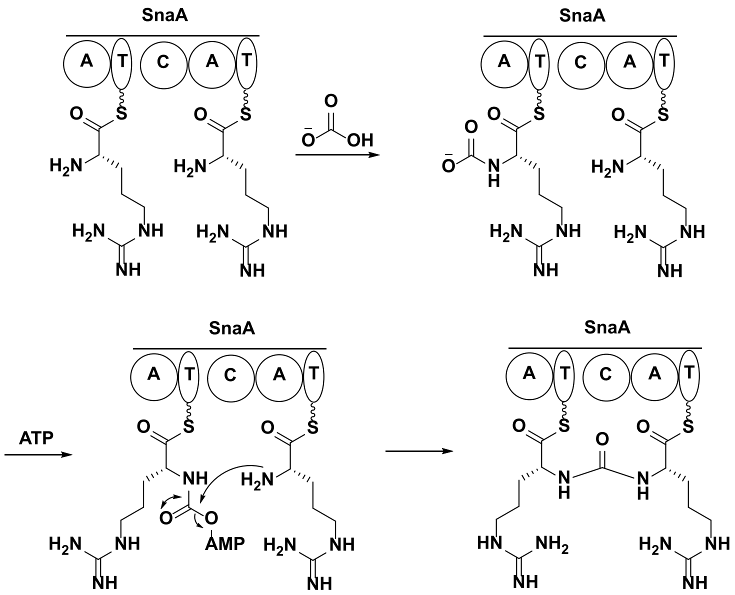


**Figure S6.** Three proposed mechanisms for ureido group formation catalyzed by SnaA. The mechanisms in panels A and B utilize the thioester linkage to activate the initially formed carbamate whereas the mechanism in panel C uses ATP for carbamate activation (shown as adenylation)

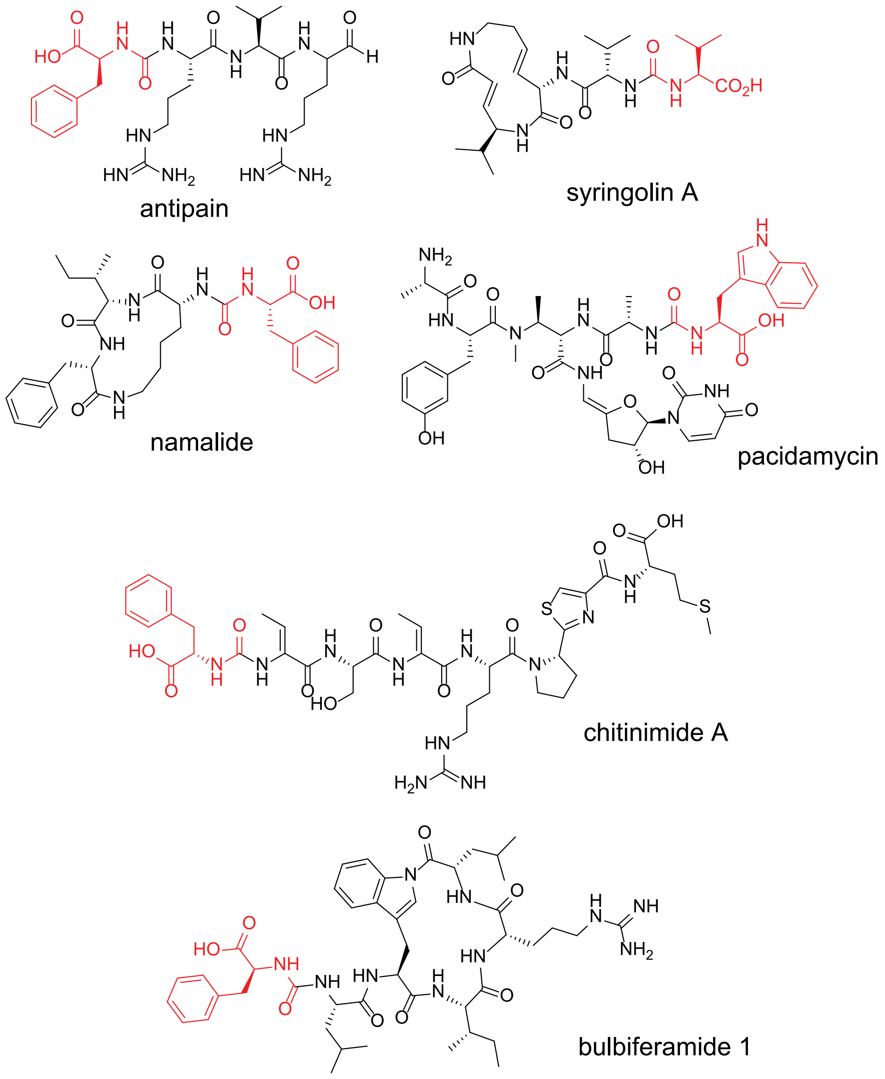


**Figure S7**. Structures of selected examples of ureido-containing nonribosomal peptides. The terminal amino acids of the ureido group have been highlighted in red. In all cases, there is only one amino acid at the terminal position of the ureido group.

**Table S1.** Primers used in this study.

| Name | Sequence |
|---|---|
| Sna1F | ATGGATCAAAACAACACCGACAAC |
| Sna1R | GATGACGGCTATGTGGGTC |
| Sna2F | CAAAGACGATCCAGTGAGGAG |
| Sna2R | ACAACTTCCAGTCCGAGC |
| Sna3F | GAACTCCTGAAGGGAGTGTC |
| Sna3R | TCTCAGTTCTTCGAGTCGC |
| Sna_OSVF | GTGGCACAGCTGGCGCGGCGACTCGAAGAACTGAGActatgttgaccgc aaactggatg |
| Sna_OSVR | TCGTTCGCGGAGTTGTCGGTGTTGTTTTGATCCATatgggtgaaccgatctc ctc |
| $SnaA_{in}$_1F | CTCTAGAAAGTATAGGAACTTCATGCATGC |
| $SnaA_{in}$_1R | ATGGTGGTGGCGGTCAGGGCATGTCCACCCAGGAGGAAC |
| $SnaA_{in}$_2F | GTTCCTCCTGGGTGGACATGCCCTGACCGCCACCACCATCGC |
| $SnaA_{in}$_2R | CAGCAGCAGGGCATGCCCGCCGAGGTCGAAG |
| $SnaA_{in}$_3F | CGGGCATGCCCTGCTGCTGGGGCATC |
| $SnaA_{in}$_3R | GTTCGATCTCGGTGAGTTCCAG |
| $SnaB_{in}$_1F | GCCAAGATCGGTTTCACCG |
| $SnaB_{in}$_1R | CTGGGTCGCCAGCAGGGCGTCGCCGCCCAGCTTG |
| $SnaB_{in}$_2F | CAAGCTGGGCGGCGACGCCCTGCTGGCGACCCAGTTG |
| $SnaB_{in}$_2R | GAAGACCCGCAGTGGCTG |
| $SnaE_{in}$_1F | CGATCGTTCGCGAGGCCCTGGGCCTCAGCGAGATGCTGCCC |
| $SnaE_{in}$_1R | CTGATCCGATTGGCACGGCGGAC |
| $SnaE_{in}$_2F | GGACGAGCGTCTGCTCCGCCATTC |
| $SnaE_{in}$_2R | GCAGGGTGTCGATCCAGGCGAAG |
| $SnaE_{in}$_3F | CGATCGGCTGCCGAGCTACATGG |
| $SnaE_{in}$_3R | GGCAGCATCTCGCTGAGGCCCAGGGCCTCGCGAACGATCGC |
| $SnaO_{in}$_1F | GCCAAGATCGGTTTCACCG |
| $SnaO_{in}$_1R | GTAGGTGTTGGCCAGGTCGTTCAGTTCGCCGGTGAC |
| $SnaO_{in}$_2F | GTCACCGGCGAACTGAACGACCTGGCCAACACCTACTG |
| $SnaO_{in}$_2R | GAAGACCCGCAGTGGCTG |
| His-TEV-SnaC_F | GGCAGCAGCCATCACCATCATCACCACGAGAACCTGTACTTCCAAA GCTTCGAAGACAATGACGGCC |
| His-TEV-SnaC_R | CCGATATCCAATTGAGATCTGCTCAAACATGGGCATCGGTC |
| SnaC_RSF_F | CATGTTTGAGCAGATCTCAATTGGATATCGG |
| SnaC_RSF_R | GGTGATGATGGTGATGGCTGCTGCCCATATGTATATCTCCTTCTTATA CTTAACTAATATACTAAGATGG |
| coSnaA1_F | CTTTAATAAGGAGATATACCATGGACCAGAATAACACCGAC |
| coSnaA1_R | CTCGTCTGTACGAATTCTTGCC |

| | |
|---|---|
| coSnaA2_F | ACAATTGCGGCAAGAATTCG |
| coSnaA2_R | AACTACTCCGCTATCAGGTAAGC |
| coSnaA3_F | GCAGGACGCTTACCTGATAG |
| coSnaA3_R | CCGGACCCGCTTTGGAAGTACAGGTTCTCCTGAGATTCTTCGCTGGACC |
| coSnaA_RSF_F | GTACTTCCAAAGCGGGTCCG |
| coSnaA_RSF_R | GGTGTTATTCTGGTCCATGGTATATCTCCTTATTAAAGTTAAACAAAATTATTTCTAC |
| coSnaAE744A_F | GCGCATCATCTGGTACATGATGGTTGG |
| coSnaAE744A_R | AACAAAGATTAACTCGTGCTCG |
| coSnaAH749A_F | GCGGATGGTTGGTCTTCTTCCCTC |
| coSnaAH749A_R | TACCAGATGATGTTCAACAAAGATTAAC |

**Table S2**. Accession IDs for proteins in the Sna BGC

| | |
|---|---|
| SnaA | ADD43706.1 |
| SnaB | ADD43707.1 |
| SnaC | ADD43708.1 |
| SnaD | ADD43709.1 |
| SnaO | ADD43710.1 |
| SnaE | ADD43711.1 |
| $SnaT_1$ | ADD43712.1 |
| $SnaT_2$ | ADD43713.1 |

**Table S3**. Accession IDs for proteins used to generate the UreaC domain multiple sequence alignment.

| Protein | MiBiG ID | NCBI ID |
|---|---|---|
| PacN | BGC0000951 | ADN26250.1 |
| NpsC | BGC0000950 | ADY76666.1 |
| Mur12 | BGC0001020 | ADZ45324.1 |
| AntC1 | BGC0001570 | OAL11435.1 |
| AntC2 | BGC0002051 | QOE83923.1 |
| RspC | BGC0000436 | EJK79842.1 |
| SylC | BGC0001047 | CAD70194.1 |
| BulbA3 | N.A. | WHI50395.1 |
| BulbA5 | N.A. | WHI48530.1 |
| PppA | BGC0002123 | QUS58939.1 |
| ChmA | BGC0002503 | QPB41096.1 |
| SnaC | N.A. | ADD43708.1 |
| ApnC1 | BGC0000301 | ABV79985.1 |
| AnaC1 | BGC0000302 | ACZ55945.1 |
| AptC1 | BGC0002512 | QNL14922.1 |
| AnaC2 | BGC0000302 | ACZ55942.1 |
| NpuC | BGC0001479 | ACC81021.1 |

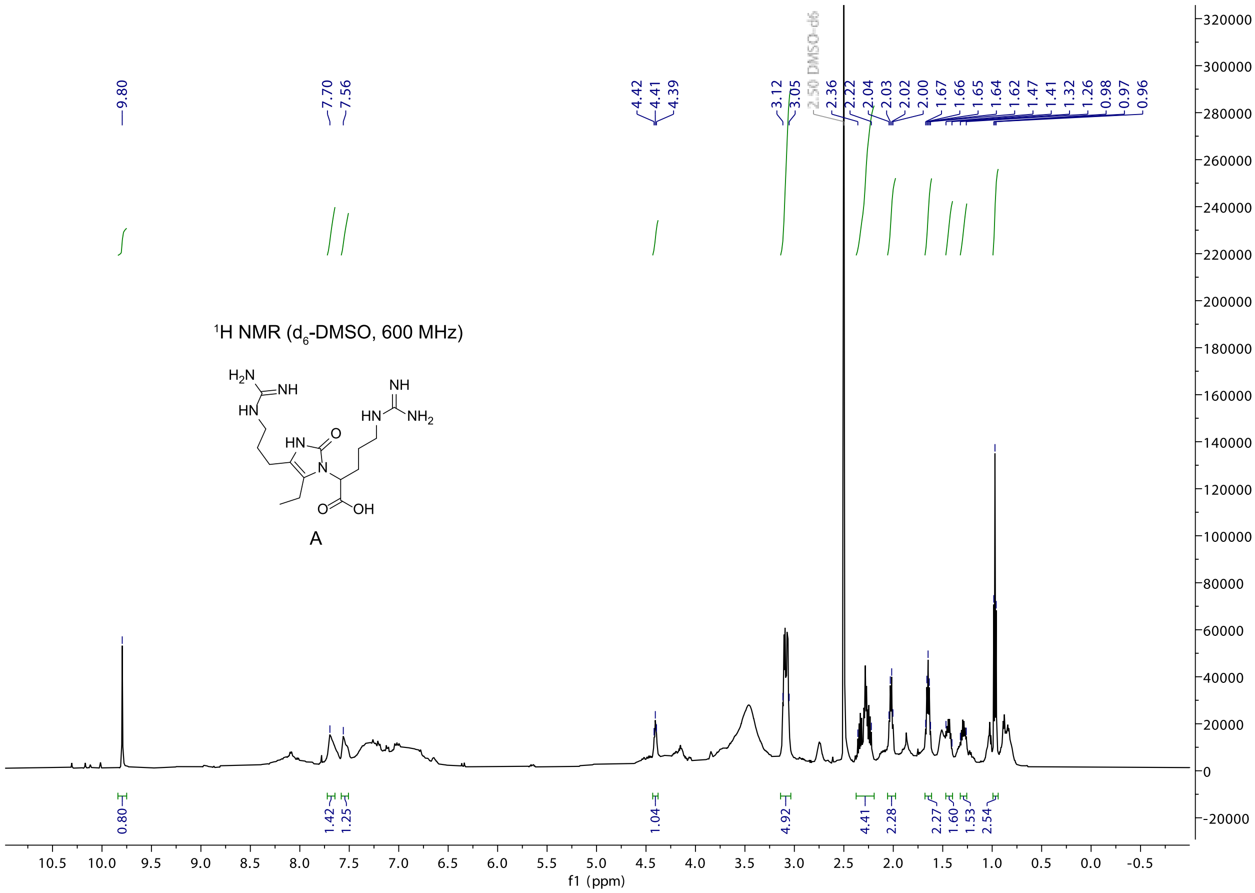


$^1$H NMR spectrum of compound A collected in $d_6$-DMSO.

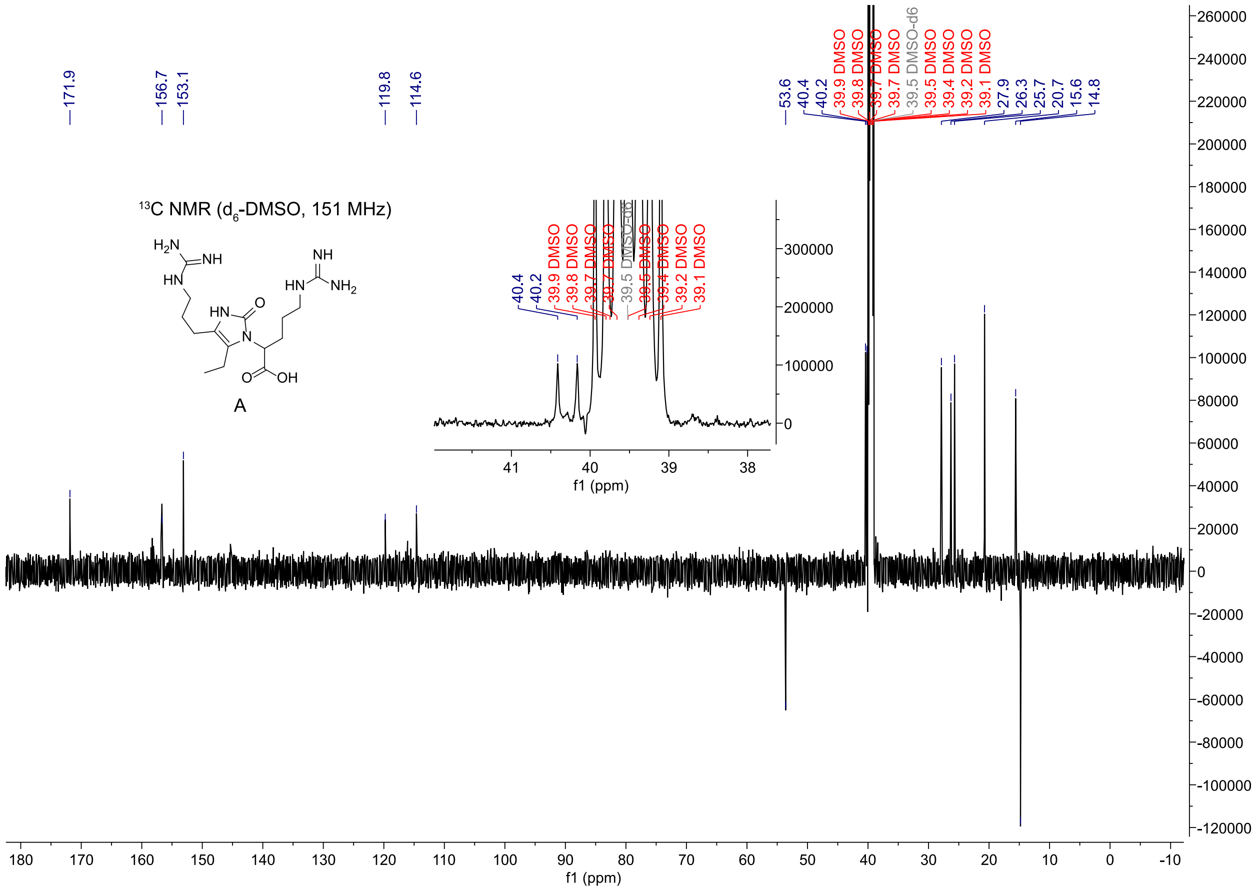


$^{13}C$ NMR spectrum of compound A collected in $d_6$-DMSO.

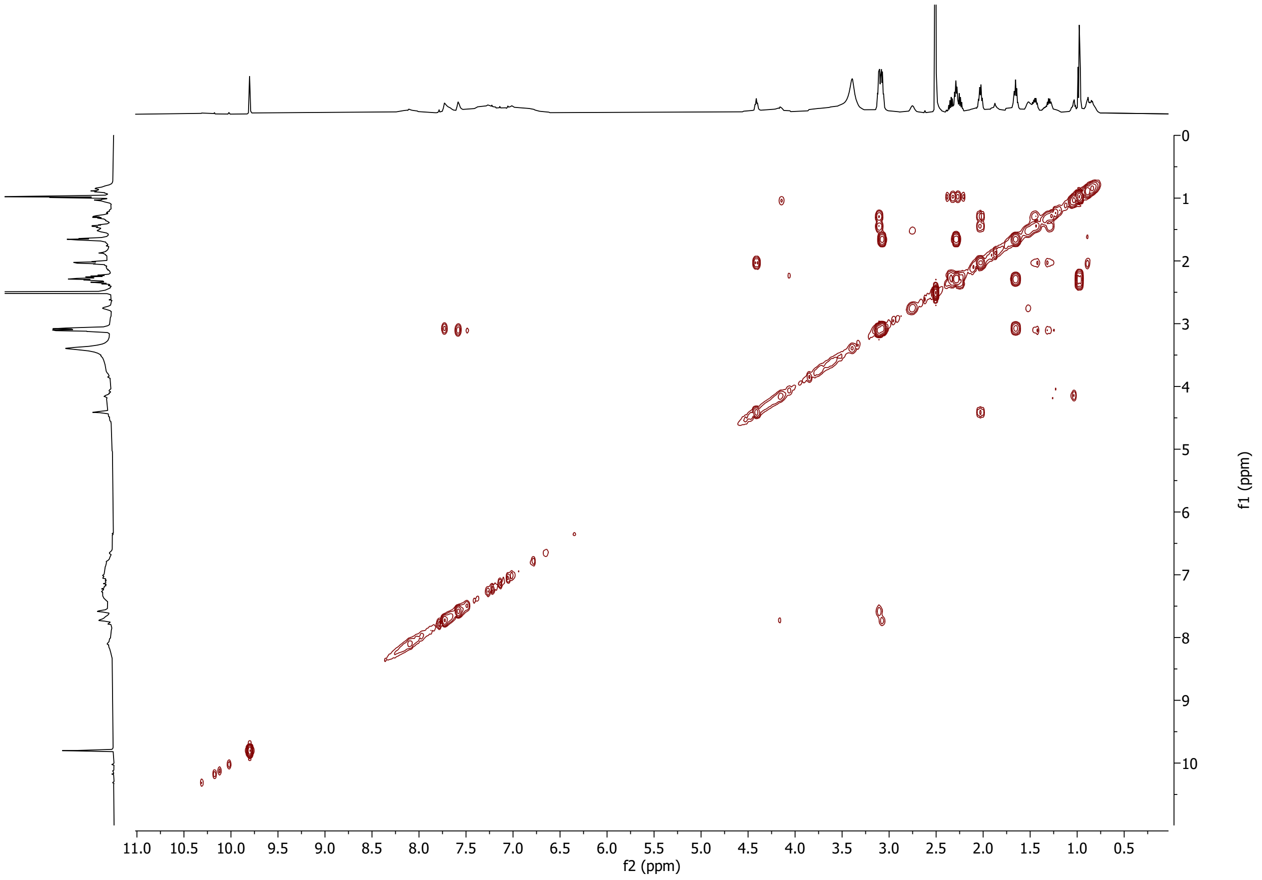


$^{1}H$-$^{1}H$ correlation spectroscopy (COSY) spectrum of compound A collected in $d_6$-DMSO.

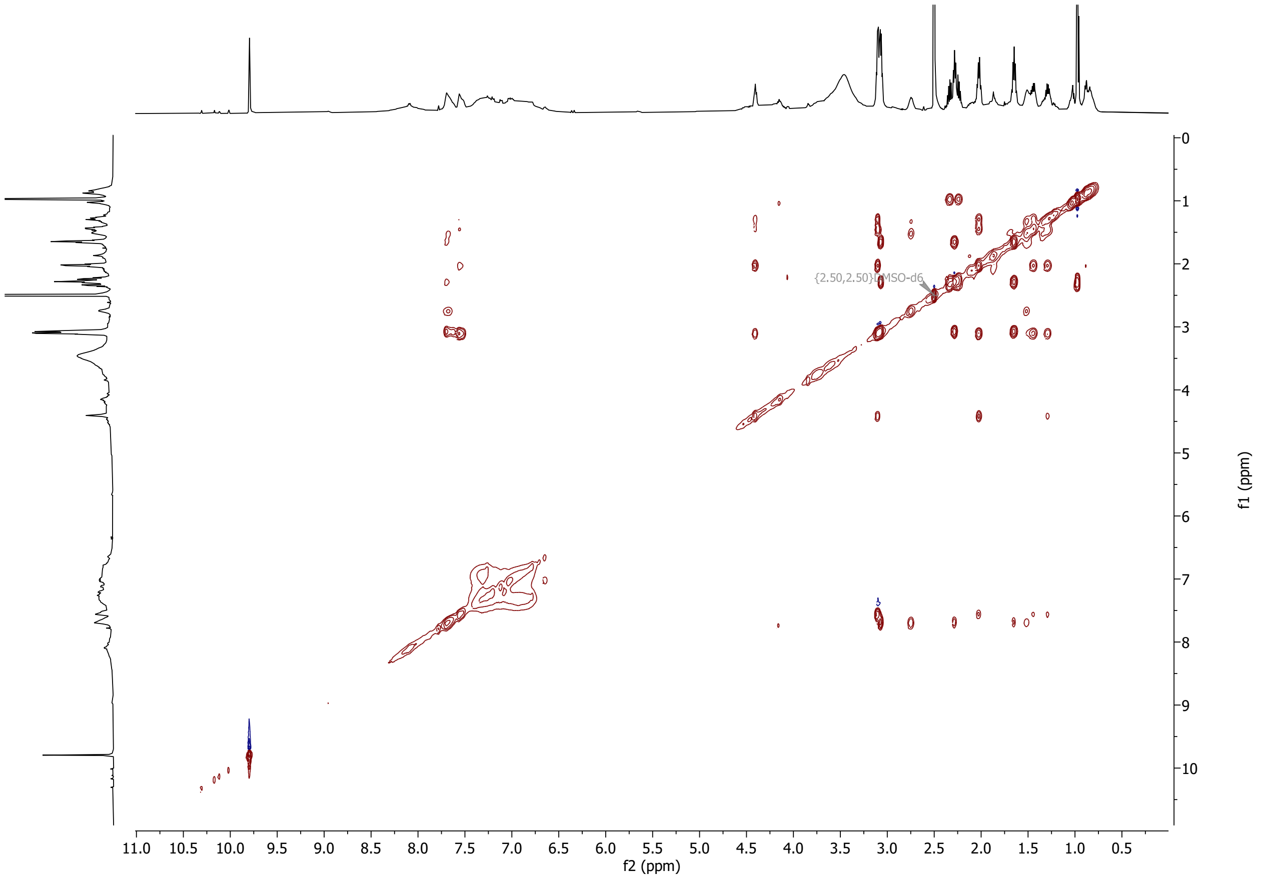


$^1$H-$^1$H total correlation spectroscopy (TOCSY) spectrum of compound A collected in $d_6$-DMSO.

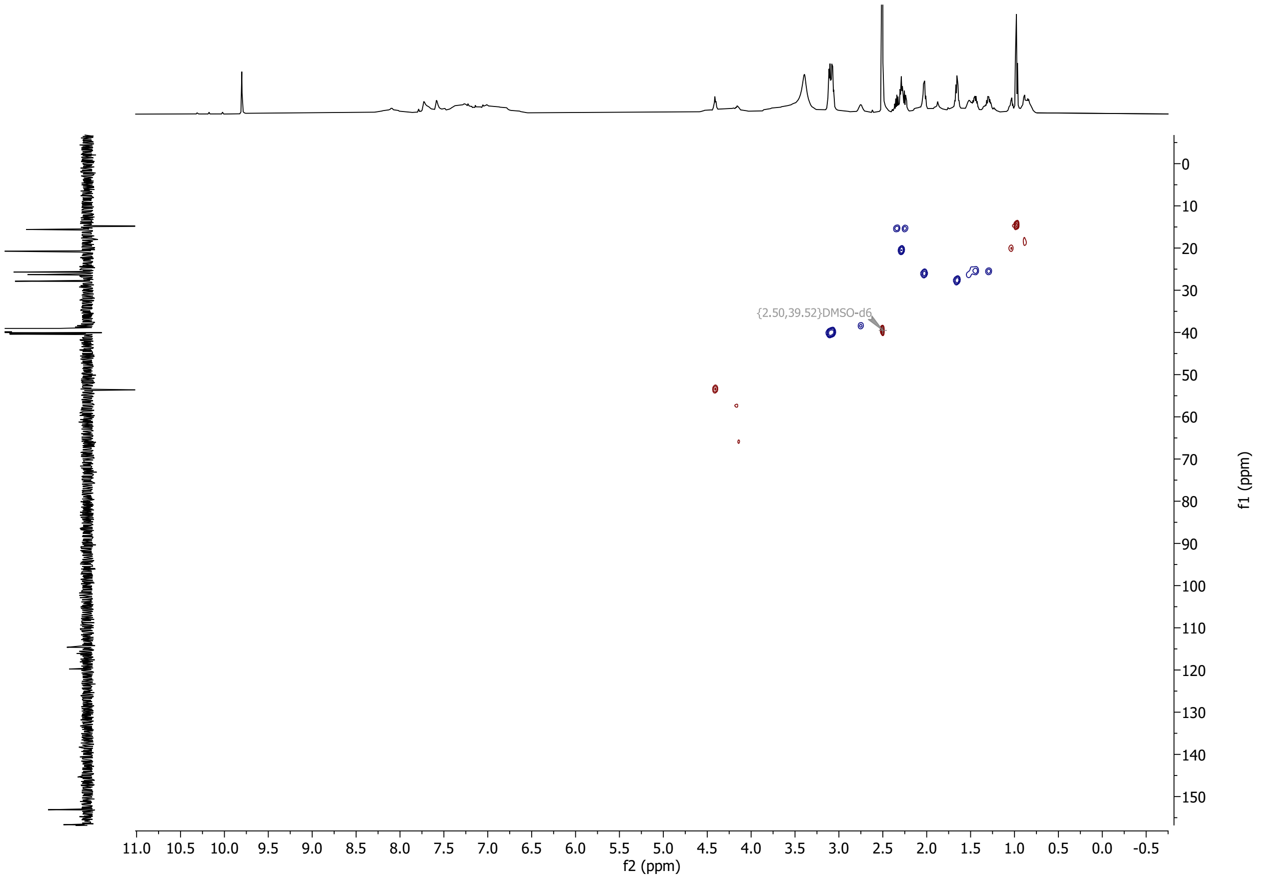


($^{1}H$,$^{13}C$) heteronuclear single quantum coherence (HSQC) spectrum of compound A collected in $d_6$-DMSO.

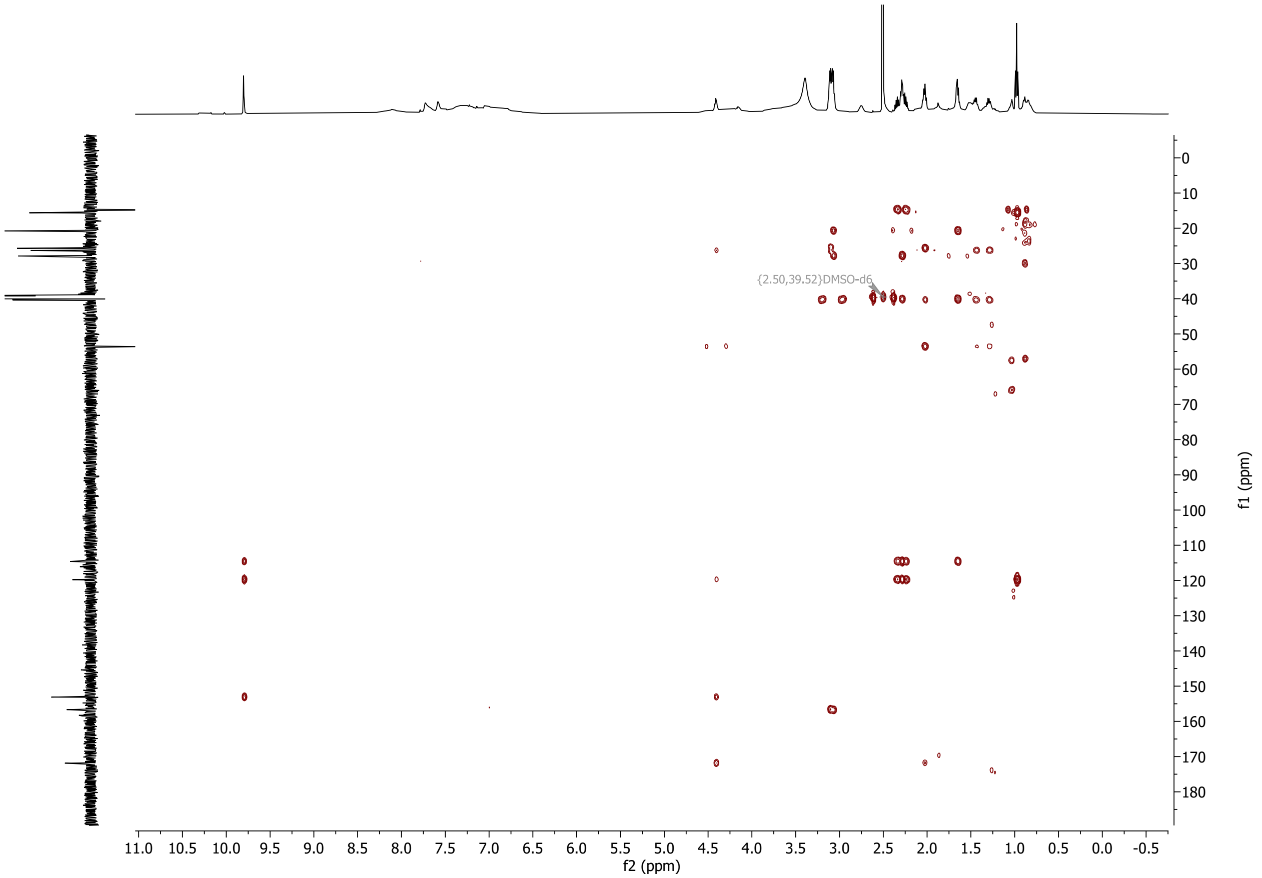


($^{1}$H,$^{13}$C) heteronuclear multiple bond correlation (HMBC) spectrum of compound A collected in $d_6$-DMSO.

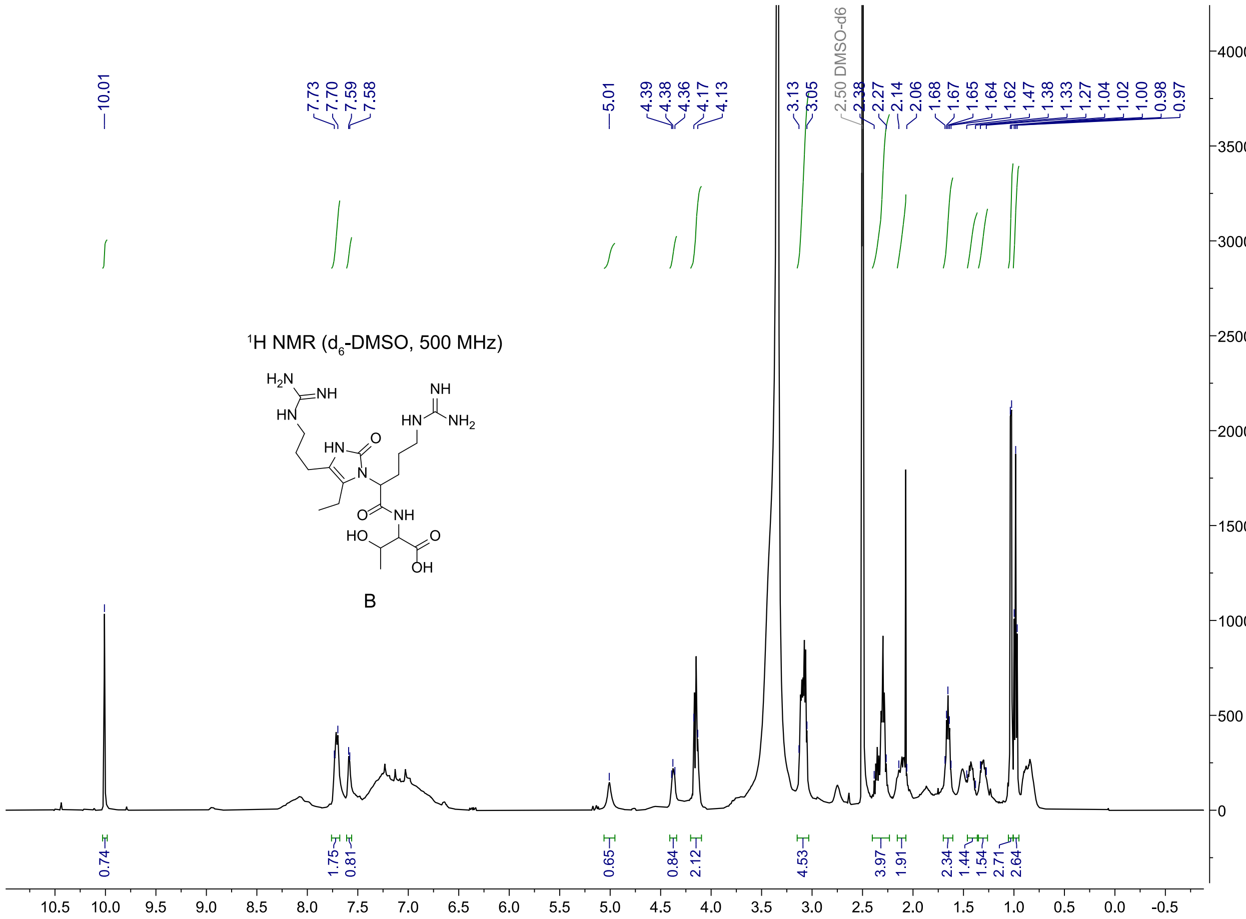


$^1$H NMR spectrum of compound B collected in $d_6$-DMSO.

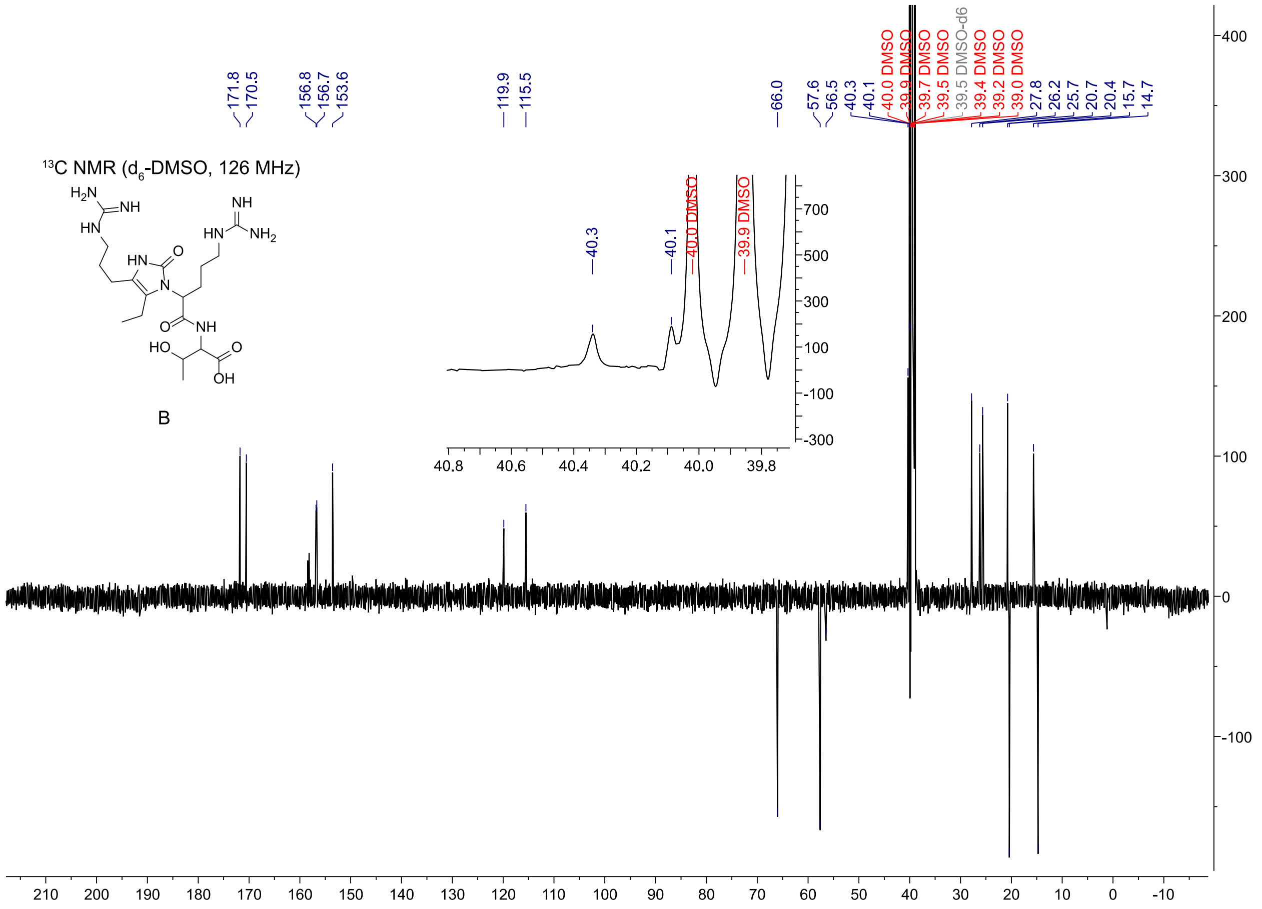


$^{13}$C NMR spectrum of compound B collected in $d_6$-DMSO.

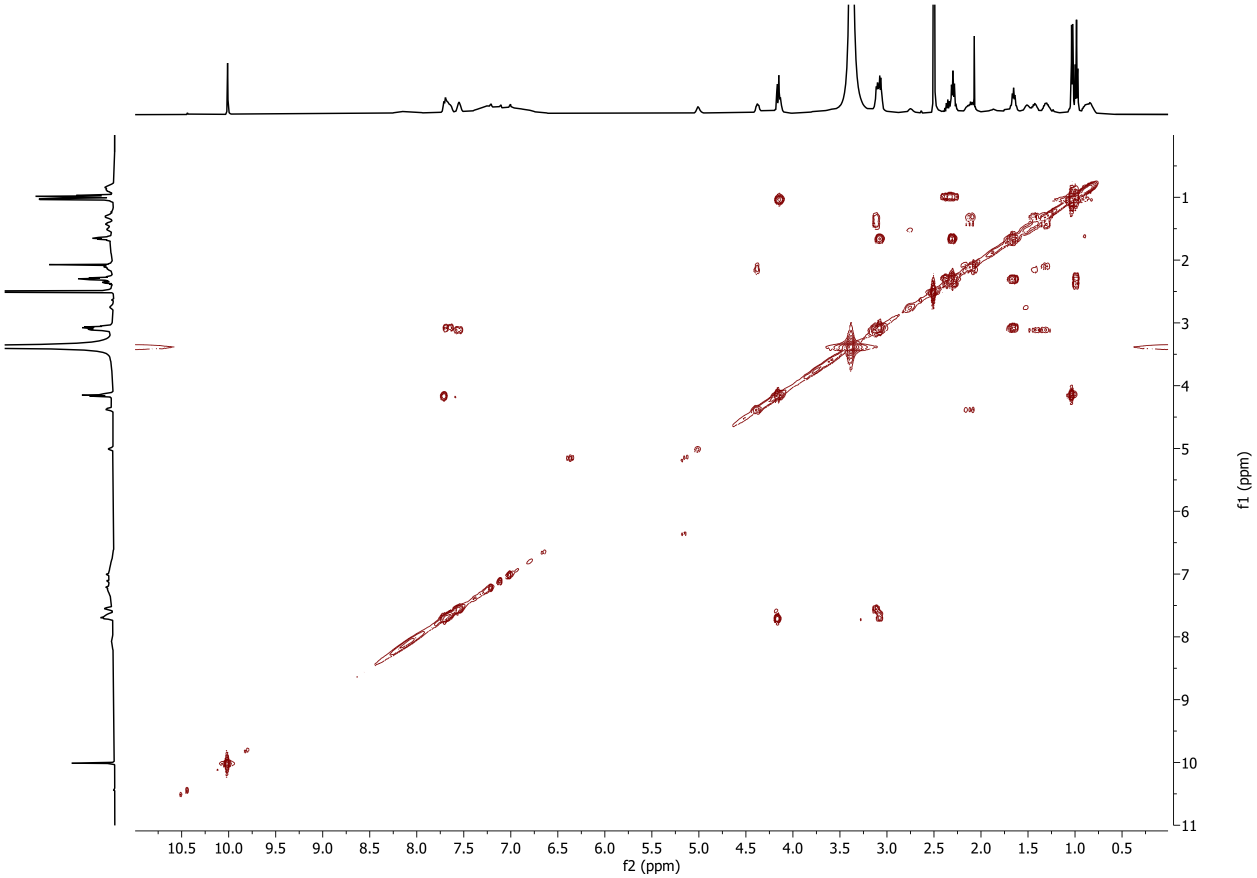


$^{1}H$-$^{1}H$ correlation spectroscopy (COSY) spectrum of compound B collected in $d_6$-DMSO.

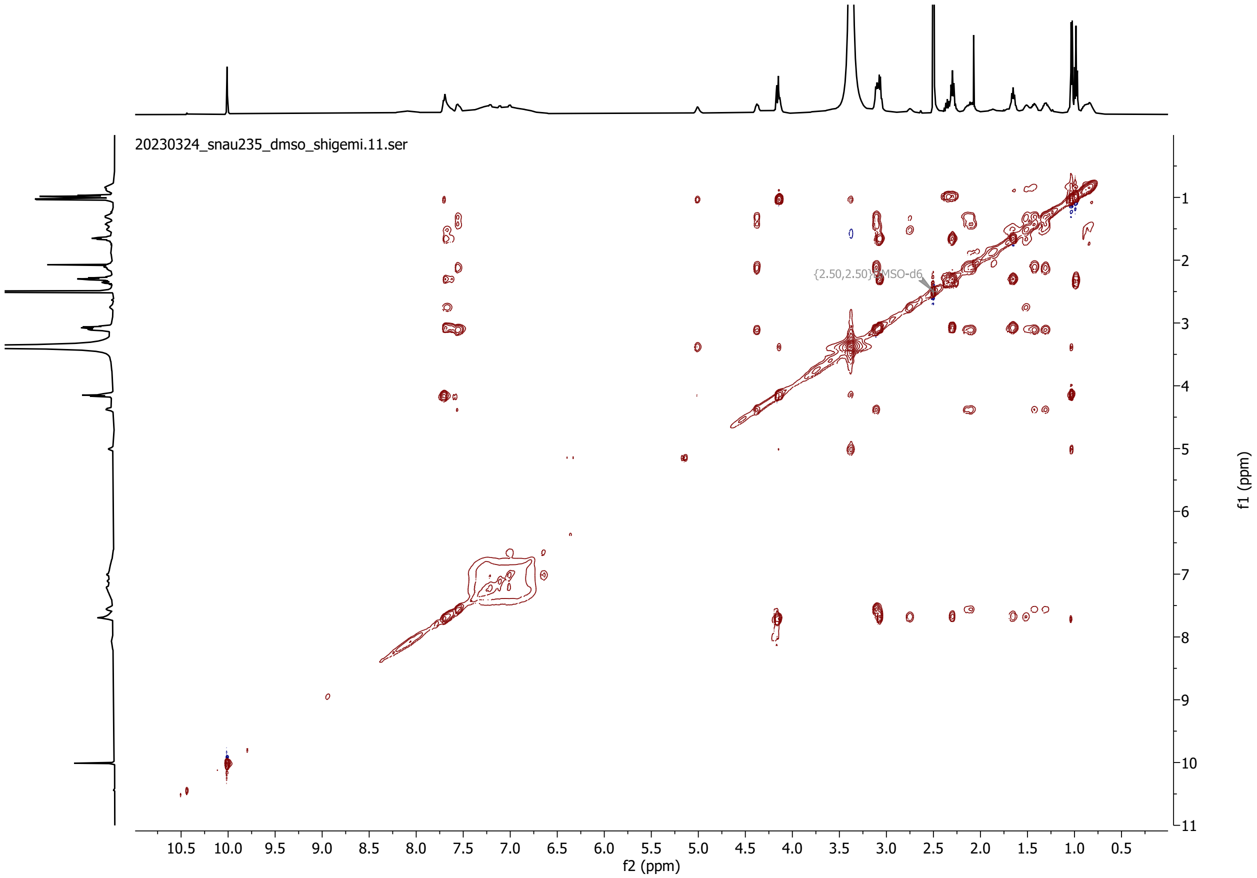


[1]H-[1]H total correlation spectroscopy (TOCSY) spectrum of compound B collected in $d_6$-DMSO.

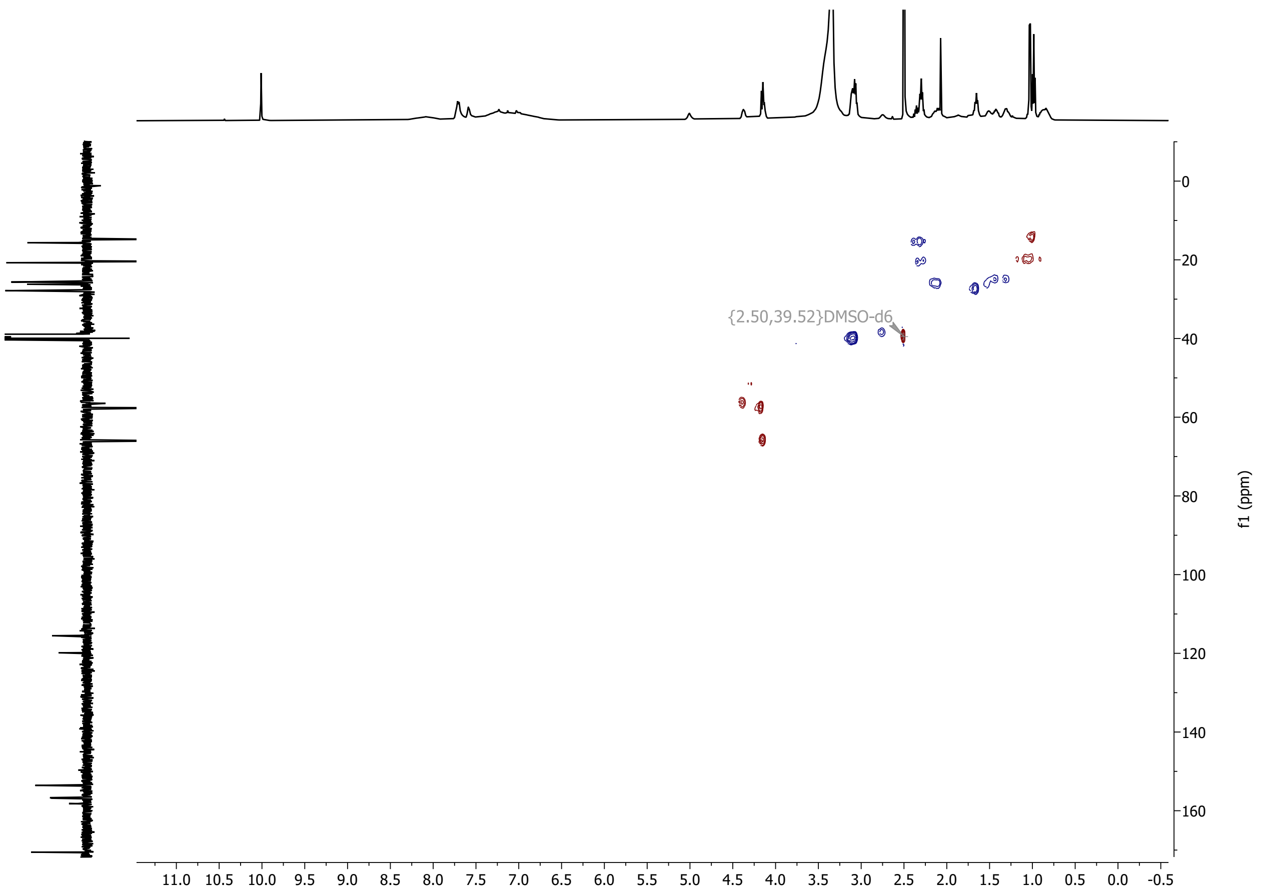


($^1$H,$^{13}$C) heteronuclear single quantum coherence (HSQC) spectrum of compound B collected in $d_6$-DMSO.

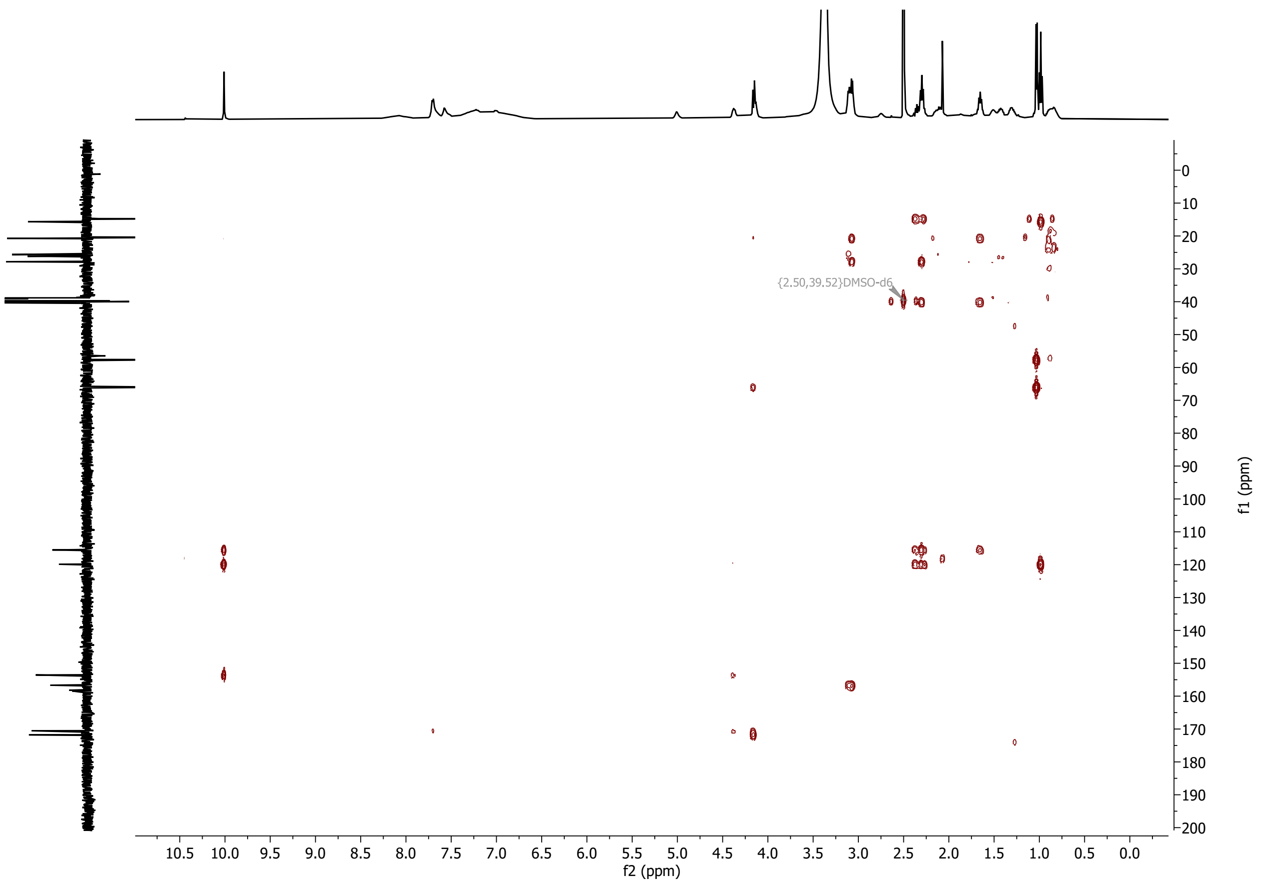


($^{1}H$,$^{13}C$) heteronuclear multiple bond correlation (HMBC) spectrum of compound B collected in $d_6$-DMSO.

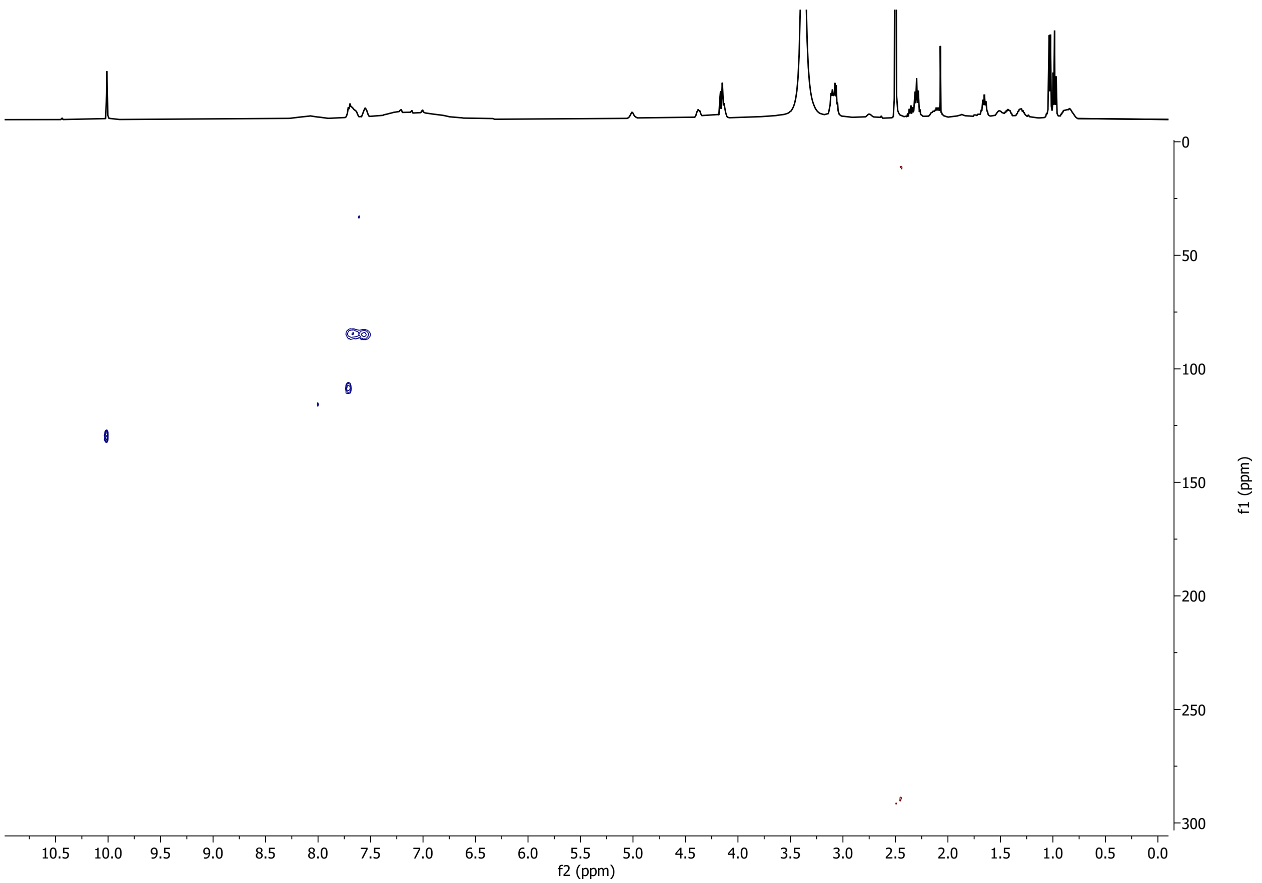


($^{1}$H,$^{15}$N) heteronuclear single quantum coherence (HSQC) spectrum of compound B collected in $d_6$-DMSO.

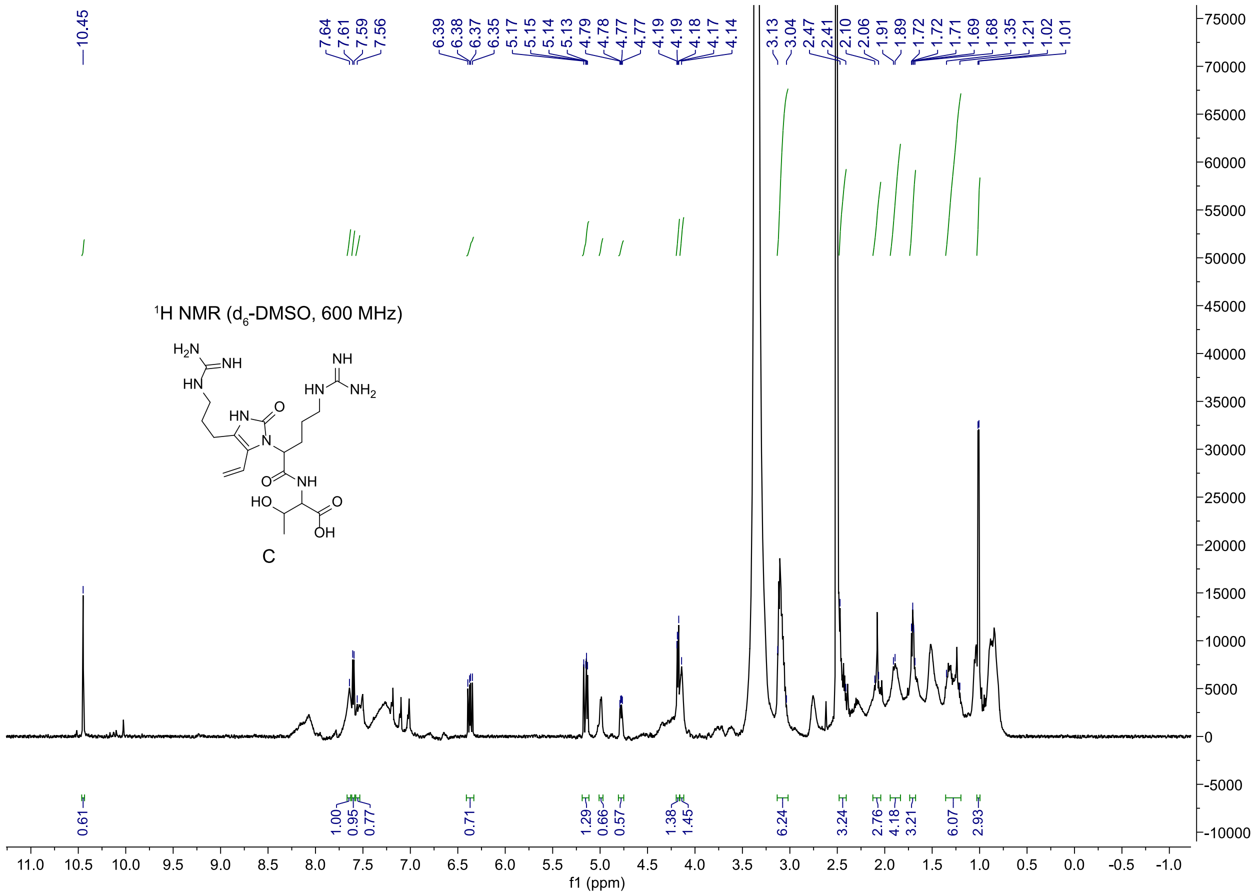


$^{1}H$ NMR spectrum of compound C collected in $d_6$-DMSO.

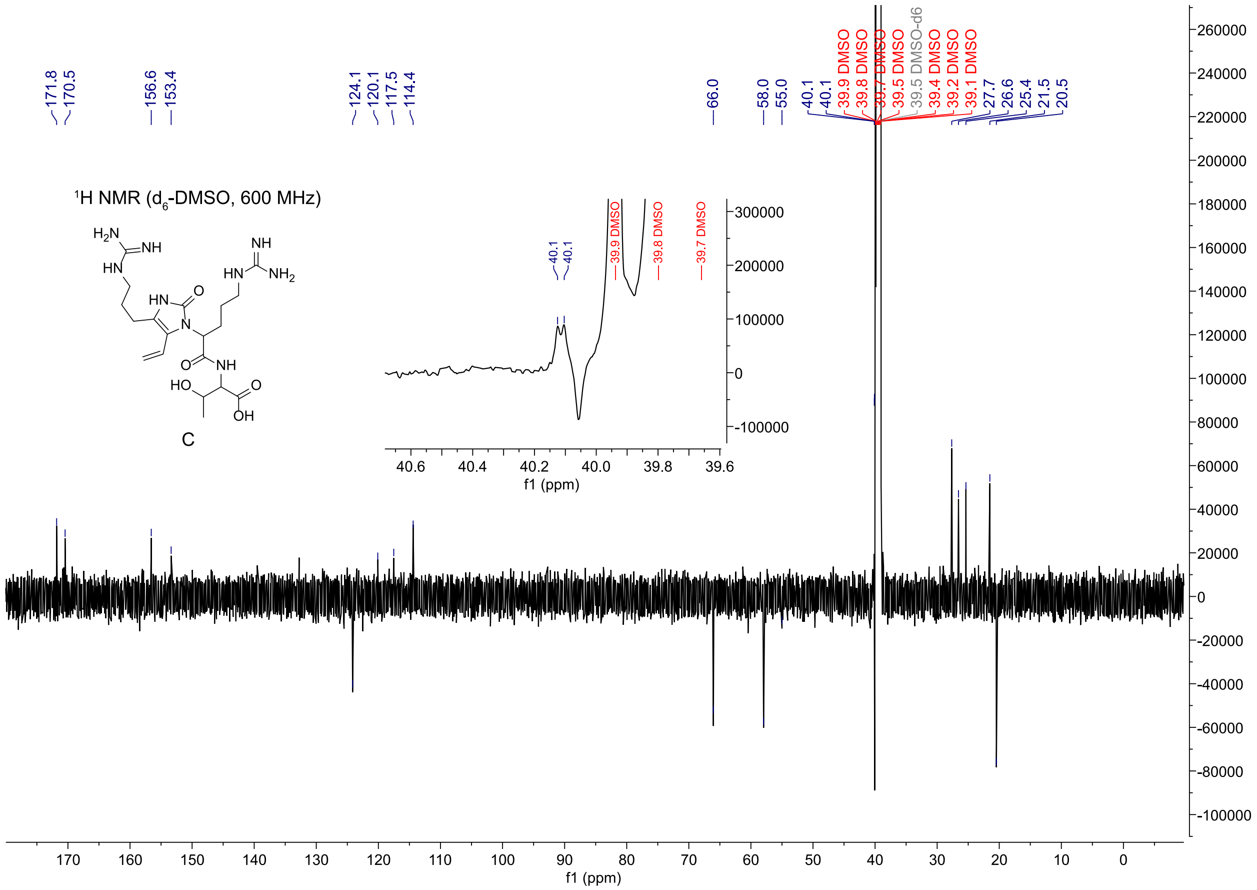


$^{13}$C NMR spectrum of compound C collected in $d_6$-DMSO.

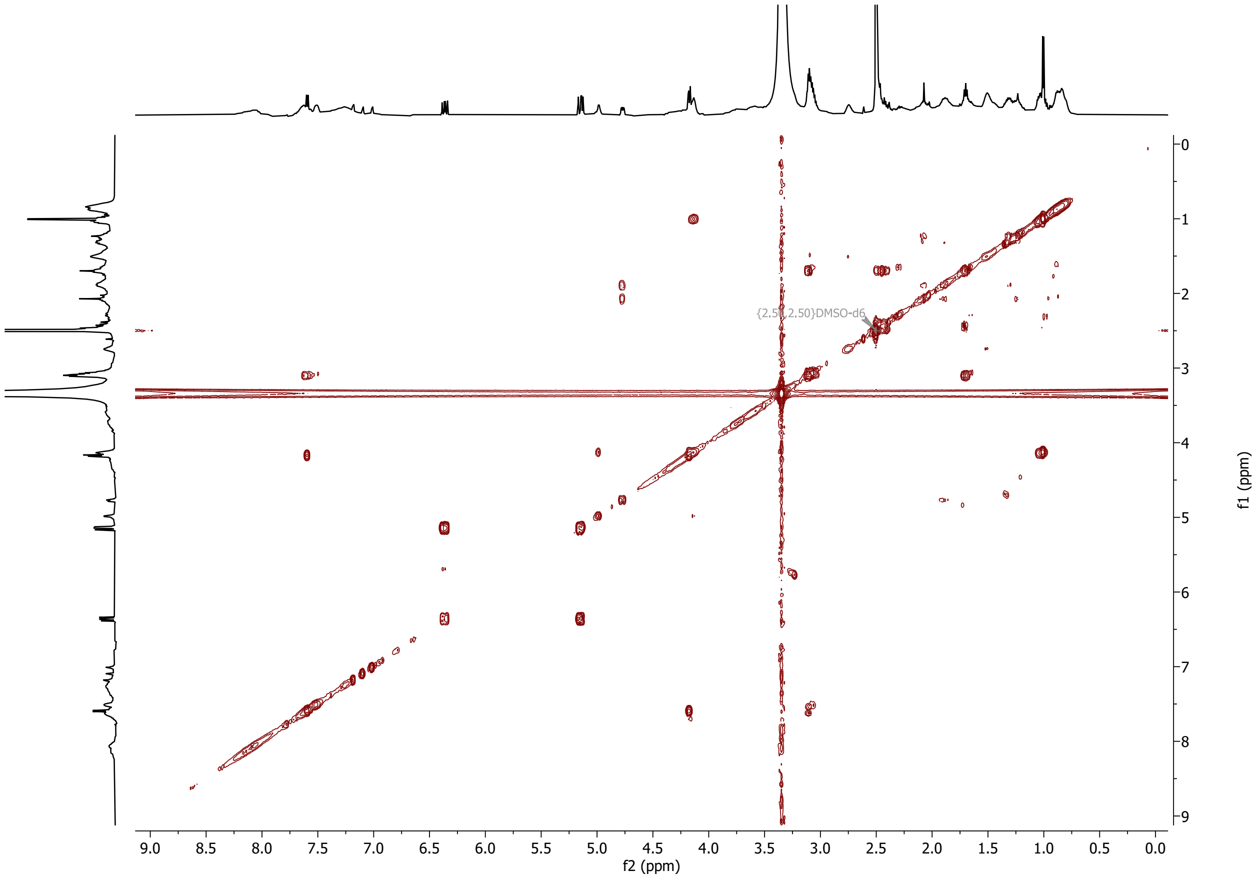


[1]H-[1]H correlation spectroscopy (COSY) spectrum of compound C collected in $d_6$-DMSO.

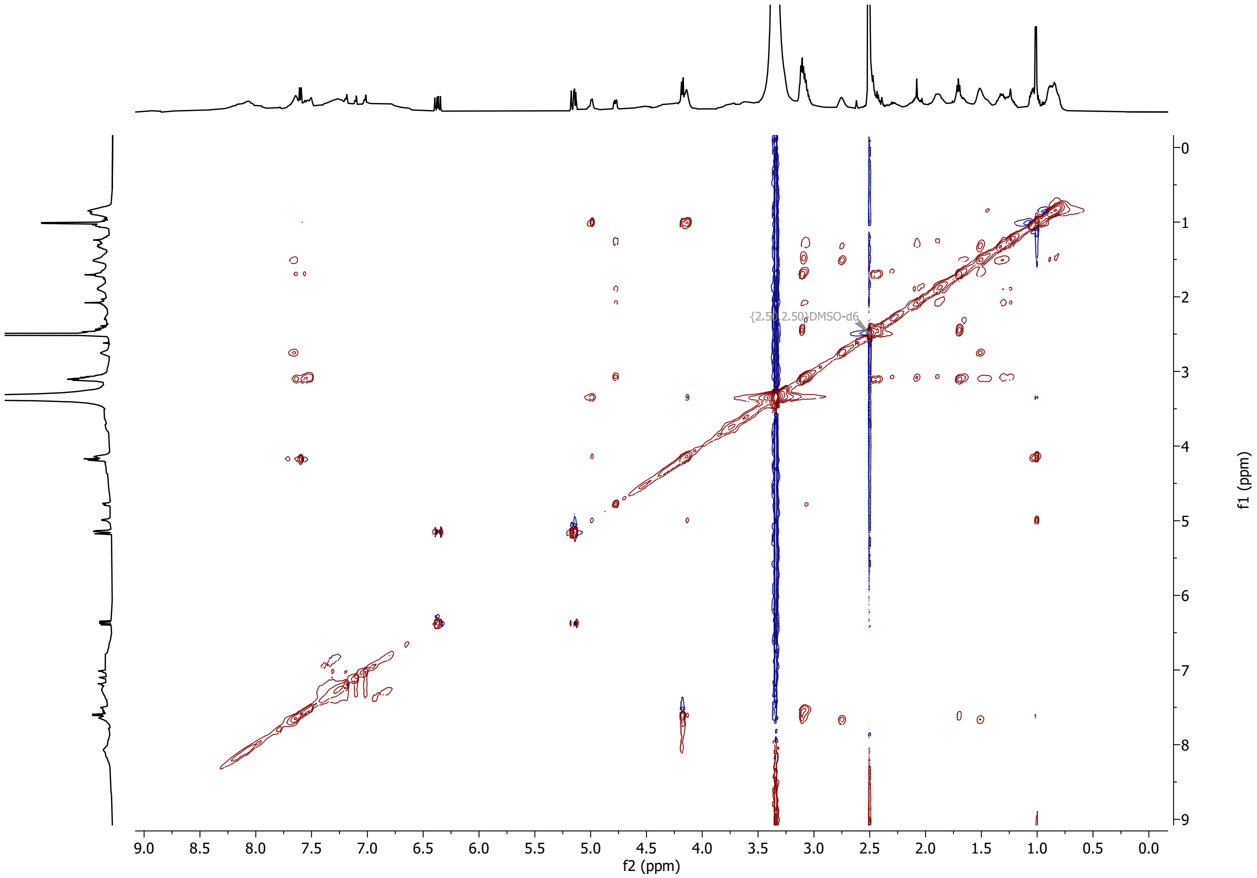


$^{1}H$-$^{1}H$ total correlation spectroscopy (TOCSY) spectrum of compound C collected in $d_6$-DMSO.

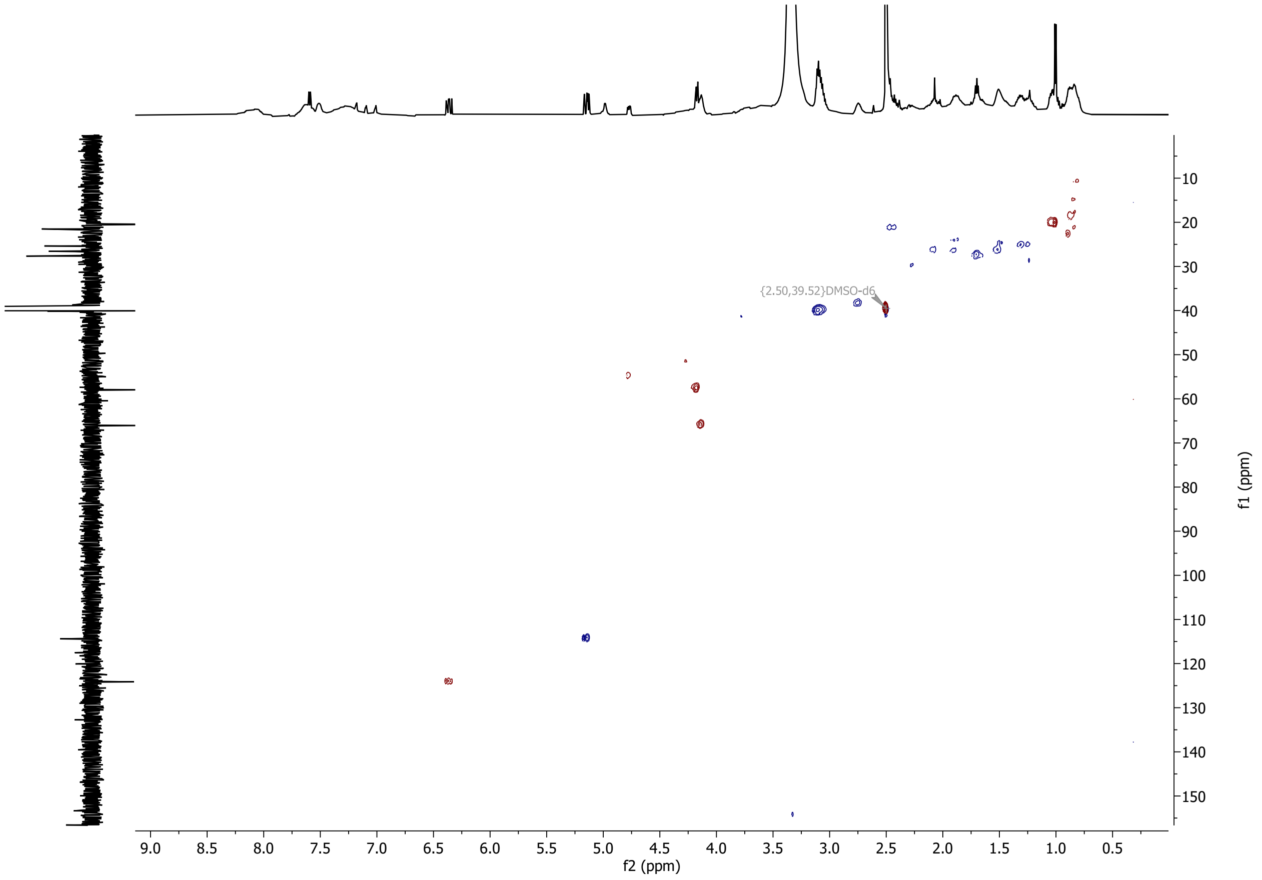


($^{1}$H,$^{13}$C) heteronuclear single quantum coherence (HSQC) spectrum of compound C collected in $d_6$-DMSO.

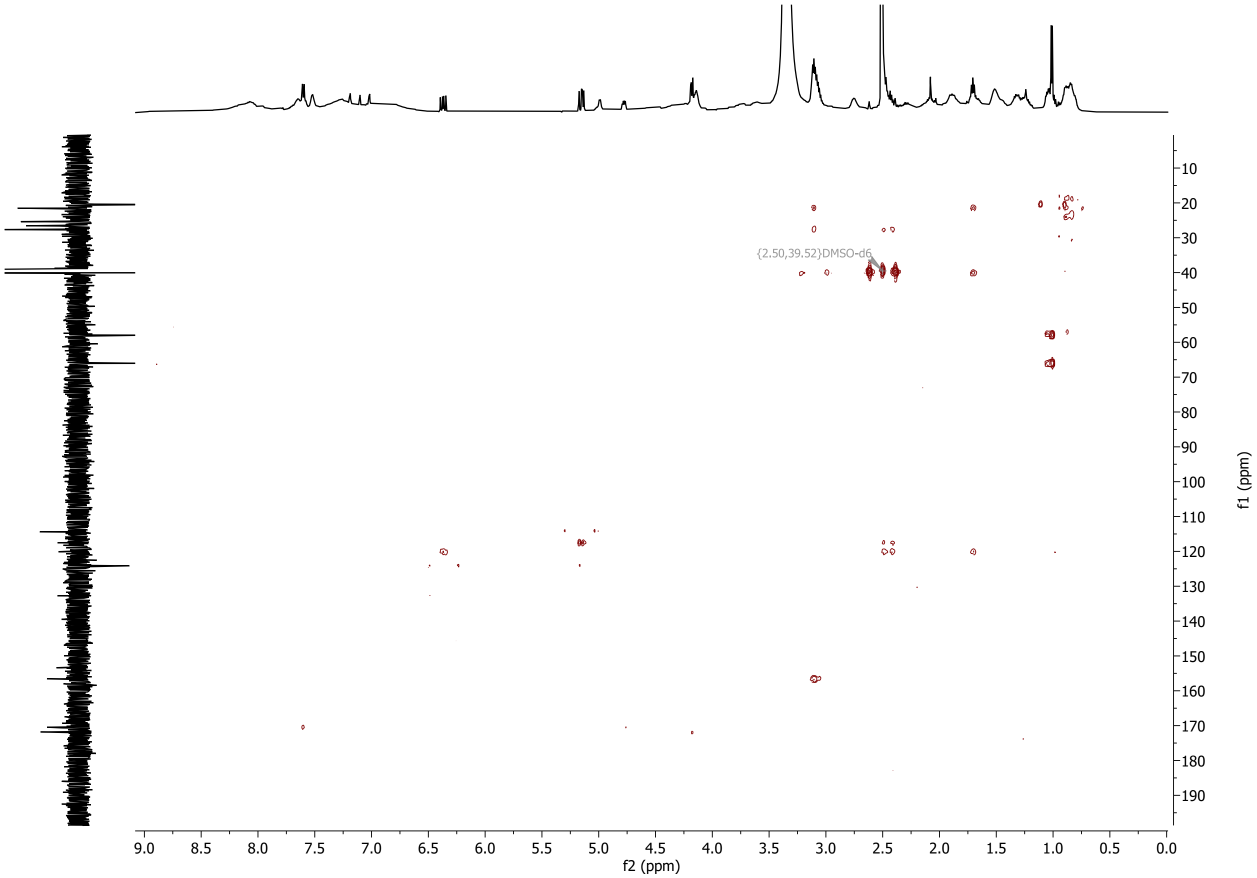


($^{1}H$,$^{13}C$) heteronuclear multiple bond correlation (HMBC) spectrum of compound C collected in $d_6$-DMSO.

### References

1. Bai, C.; Zhang, Y.; Zhao, X.; Hu, Y.; Xiang, S.; Miao, J.; Lou, C.; Zhang, L., Exploiting a precise design of universal synthetic modular regulatory elements to unlock the microbial natural products in *Streptomyces*. *Proc. Natl. Acad. Sci. U. S. A.* **2015,** *112* (39), 12181-6.

2. Gibson, D. G.; Young, L.; Chuang, R. Y.; Venter, J. C.; Hutchison, C. A., 3rd; Smith, H. O., Enzymatic assembly of DNA molecules up to several hundred kilobases. *Nat. Methods* **2009,** *6* (5), 343-5.

3. Kieser, T.; Bibb, M. J.; Buttner, M. J.; Chater, K. F.; Hopwood, D. A., Practical Streptomyces Genetics. **2000**.

4. Kim, W.; Hwang, S.; Lee, N.; Lee, Y.; Cho, S.; Palsson, B.; Cho, B. K., Transcriptome and translatome profiles of Streptomyces species in different growth phases. *Sci. Data* **2020,** *7* (1), 138.

5. Chambers, M. C.; Maclean, B.; Burke, R.; Amodei, D.; Ruderman, D. L.; Neumann, S.; Gatto, L.; Fischer, B.; Pratt, B.; Egertson, J.; Hoff, K.; Kessner, D.; Tasman, N.; Shulman, N.; Frewen, B.; Baker, T. A.; Brusniak, M. Y.; Paulse, C.; Creasy, D.; Flashner, L.; Kani, K.; Moulding, C.; Seymour, S. L.; Nuwaysir, L. M.; Lefebvre, B.; Kuhlmann, F.; Roark, J.; Rainer, P.; Detlev, S.; Hemenway, T.; Huhmer, A.; Langridge, J.; Connolly, B.; Chadick, T.; Holly, K.; Eckels, J.; Deutsch, E. W.; Moritz, R. L.; Katz, J. E.; Agus, D. B.; MacCoss, M.; Tabb, D. L.; Mallick, P., A cross-platform toolkit for mass spectrometry and proteomics. *Nat. Biotechnol.* **2012,** *30* (10), 918-20.

6. Smith, C. A.; Want, E. J.; O'Maille, G.; Abagyan, R.; Siuzdak, G., XCMS: processing mass spectrometry data for metabolite profiling using nonlinear peak alignment, matching, and identification. *Anal. Chem.* **2006,** *78* (3), 779-87.

7. Maxson, T.; Tietz, J. I.; Hudson, G. A.; Guo, X. R.; Tai, H. C.; Mitchell, D. A., targeting reactive carbonyls for identifying natural products and their biosynthetic origins. *J. Am. Chem. Soc.* **2016,** *138* (46), 15157-15166.

8. Iorio, M.; Davatgarbenam, S.; Serina, S.; Criscenzo, P.; Zdouc, M. M.; Simone, M.; Maffioli, S. I.; Ebright, R. H.; Donadio, S.; Sosio, M., Blocks in the pseudouridimycin pathway unlock hidden metabolites in the *Streptomyces* producer strain. *Scientific reports* **2021,** *11* (1), 5827.

9. Cui, Z.; Nguyen, H.; Bhardwaj, M.; Wang, X.; Büschleb, M.; Lemke, A.; Schütz, C.; Rohrbacher, C.; Junghanns, P.; Koppermann, S.; Ducho, C.; Thorson, J. S.; Van Lanen, S. G., Enzymatic C(β)-H functionalization of L-Arg and L-Leu in nonribosomally derived peptidyl natural products: a tale of two oxidoreductases. *J. Am. Chem. Soc.* **2021,** *143* (46), 19425-19437.

10. Christiansen, G.; Philmus, B.; Hemscheidt, T.; Kurmayer, R., Genetic variation of adenylation domains of the anabaenopeptin synthesis operon and evolution of substrate promiscuity. *J. Bacteriol.* **2011,** *193* (15), 3822-31.

11. Rouhiainen, L.; Jokela, J.; Fewer, D. P.; Urmann, M.; Sivonen, K., Two alternative starter modules for the non-ribosomal biosynthesis of specific anabaenopeptin variants in *Anabaena* (Cyanobacteria). *Chem. Biol.* **2010,** *17* (3), 265-73.

12. Greunke, C.; Duell, E. R.; D'Agostino, P. M.; Glöckle, A.; Lamm, K.; Gulder, T. A. M., Direct Pathway Cloning (DiPaC) to unlock natural product biosynthetic potential. *Metab. Eng.* **2018,** *47*, 334-345.

13. Koketsu, K.; Mitsuhashi, S.; Tabata, K., Identification of homophenylalanine biosynthetic genes from the cyanobacterium *Nostoc punctiforme* PCC73102 and application to its microbial production by *Escherichia coli*. *Appl. Environ. Microbiol.* **2013,** *79* (7), 2201-8.

14. Saha, S.; Esposito, G.; Urajová, P.; Mareš, J.; Ewe, D.; Caso, A.; Macho, M.; Delawská, K.; Kust, A.; Hrouzek, P.; Juráň, J.; Costantino, V.; Saurav, K., Discovery of unusual cyanobacterial tryptophan-containing anabaenopeptins by MS/MS-based molecular networking. *Molecules* **2020,** *25* (17).

15. Dudnik, A.; Bigler, L.; Dudler, R., Production of proteasome inhibitor syringolin A by the endophyte *Rhizobium* sp. strain AP16. *Appl. Environ. Microbiol.* **2014,** *80* (12), 3741-8.

16. Amrein, H.; Makart, S.; Granado, J.; Shakya, R.; Schneider-Pokorny, J.; Dudler, R., Functional analysis of genes involved in the synthesis of syringolin A by *Pseudomonas syringae* pv. *syringae* B301 D-R. *Mol. Plant. Microbe. Interact.* **2004,** *17* (1), 90-7.
17. Zhang, W.; Ostash, B.; Walsh, C. T., Identification of the biosynthetic gene cluster for the pacidamycin group of peptidyl nucleoside antibiotics. *Proc. Natl. Acad. Sci. U. S. A.* **2010,** *107* (39), 16828-33.
18. Kaysser, L.; Tang, X.; Wemakor, E.; Sedding, K.; Hennig, S.; Siebenberg, S.; Gust, B., Identification of a napsamycin biosynthesis gene cluster by genome mining. *ChemBioChem* **2011,** *12* (3), 477-87.
19. Tang, X.; Gross, M.; Xie, Y.; Kulik, A.; Gust, B., Identification of mureidomycin analogues and functional analysis of an N-acetyltransferase in napsamycin biosynthesis. *ChemBioChem* **2013,** *14* (17), 2248-55.
20. Cheng, L.; Chen, W.; Zhai, L.; Xu, D.; Huang, T.; Lin, S.; Zhou, X.; Deng, Z., Identification of the gene cluster involved in muraymycin biosynthesis from *Streptomyces* sp. NRRL 30471. *Mol. Biosyst.* **2011,** *7* (3), 920-7.
21. Liu, J.; Zhou, H.; Yang, Z.; Wang, X.; Chen, H.; Zhong, L.; Zheng, W.; Niu, W.; Wang, S.; Ren, X.; Zhong, G.; Wang, Y.; Ding, X.; Müller, R.; Zhang, Y.; Bian, X., Rational construction of genome-reduced Burkholderiales chassis facilitates efficient heterologous production of natural products from proteobacteria. *Nat. Commun.* **2021,** *12* (1), 4347.
22. Ióca, L. P.; Dai, Y.; Kunakom, S.; Diaz-Espinosa, J.; Krunic, A.; Crnkovic, C. M.; Orjala, J.; Sanchez, L. M.; Ferreira, A. G.; Berlinck, R. G. S.; Eustáquio, A. S., A family of nonribosomal peptides modulate collective behavior in *Pseudovibrio* bacteria isolated from marine sponges. *Angew. Chem. Int. Ed. Engl.* **2021,** *60* (29), 15891-15898.
23. Zhong, W.; Deutsch, J. M.; Yi, D.; Abrahamse, N. H.; Mohanty, I.; Moore, S. G.; McShan, A. C.; Garg, N.; Agarwal, V., Discovery and biosynthesis of ureidopeptide natural products macrocyclized via indole N-acylation in marine *Microbulbifer spp.* Bacteria. *ChemBioChem* **2023,** *24* (12), e202300190.
24. Zhong, W.; Aiosa, N.; Deutsch, J. M.; Garg, N.; Agarwal, V., Pseudobulbiferamides: plasmid-encoded ureidopeptide natural products with biosynthetic gene clusters shared among marine bacteria of different genera. *J. Nat. Prod.* **2023,** *86* (10), 2414-2420.
25. Imker, H. J.; Walsh, C. T.; Wuest, W. M., SylC catalyzes ureido-bond formation during biosynthesis of the proteasome inhibitor syringolin A. *J. Am. Chem. Soc.* **2009,** *131* (51), 18263-5.
